# Gene co-expression network reveals highly conserved, well-regulated anti-ageing mechanisms in old ant queens

**DOI:** 10.1101/2021.02.14.431190

**Authors:** Mark C. Harrison, Luisa M. Jaimes Niño, Marisa Almeida Rodrigues, Judith Ryll, Thomas Flatt, Jan Oettler, Erich Bornberg-Bauer

## Abstract

Evolutionary theories of ageing predict a reduction in selection efficiency with age, a so-called ‘selection shadow’, due to extrinsic mortality decreasing effective population size with age. Classic symptoms of ageing include a deterioration in transcriptional regulation and protein homeostasis. Understanding how ant queens defy the trade-off between fecundity and lifespan remains a major challenge for the evolutionary theory of ageing. It has often been discussed that the low extrinsic mortality of ant queens, that are generally well protected within the nest by workers and soldiers, should reduce the selection shadow acting on old queens. We tested this by comparing strength of selection acting on genes upregulated in young and old queens of the ant, *Cardiocondyla obscurior*. In support of a reduced selection shadow, we find old-biased genes to be under strong purifying selection. We also analysed a gene co-expression network (GCN) with the aim to detect signs of ageing in the form of deteriorating regulation and proteostasis. We find no evidence for ageing. In fact, we detect higher connectivity in old queens indicating increased transcriptional regulation with age. Within the GCN, we discover five highly correlated modules that are upregulated with age. These old-biased modules regulate several anti-ageing mechanisms such as maintenance of proteostasis, transcriptional regulation and stress response. We observe stronger purifying selection on central hub genes of these old-biased modules compared to young-biased modules. These results indicate a lack of transcriptional ageing in old *C. obscurior* queens possibly facilitated by strong selection at old age and well-regulated anti-ageing mechanisms.

**Significance Statement:** Understanding the exceptional longevity of ant queens and how they defy the trade-off between fecundity and lifespan remains a major challenge for the evolutionary theory and molecular biology of ageing. In this study we offer several clues as to how this occurs on a molecular level in *C. obscurior* queens. Specifically, we believe a reduction in the selection shadow due to low extrinsic mortality, has allowed the evolution of well-regulated anti-ageing mechanisms. Consequently, we suggest several promising starting points for future research into the poorly understood phenomenon of extreme longevity in ant queens. Making progress in this field will not only allow us to better understand longevity and fertility in social insects but may also offer interesting research strategies for human ageing.

## Introduction

Ageing, the progressive decline of physiological function with age, and thus of survival and fertility, is common to most multicellular species (Jones et al., 2014). Extensive genetic and molecular studies have illuminated several proximate mechanisms involved in the ageing process, allowing us to better understand *how* we age. The majority of these “hallmarks of ageing” can be attributed to the accumulation of cellular damage (López-Otín et al., 2013; Gems and Partridge, 2013) and an overall deterioration of regulation (Frenk and Houseley, 2018). One important hallmark of ageing, the loss of protein homeostasis, is caused by a reduction in quality control mechanisms such as chaperones that support correct folding and structure of proteins, as well as proteolytic pathways that ensure the removal of misfolded peptides (Koga et al., 2011; Rubinsztein et al., 2011; Tomaru et al., 2012; Calderwood et al., 2009; López-Otín et al., 2013). The result is an accumulation of toxic, misfolded proteins and an inefficient replenishment of correctly functioning proteins. Further hallmarks of ageing include deleterious changes in terms of cell-cycle (a cessation of cellular replication), intercellular communication, nutrient sensing and epigenetic regulation (López-Otín et al., 2013), as well as a downregulation of mitochondrial and protein synthesis genes (Frenk and Houseley, 2018). Importantly, the ageing process is often accompanied by a dysregulation of transcription (Frenk and Houseley, 2018).

Several classic evolutionary theories of ageing aim to explain *why* organisms age (Kirkwood and Austad, 2000; Flatt and Partridge, 2018). These theories generally describe a reduction in selection efficiency with increasing age because the number of surviving individuals decreases due to extrinsic mortality. In the mutation accumulation theory, this ‘selection shadow’ leads to an accumulation of mutations which have a deleterious effect later in life (Kirkwood and Austad, 2000; Flatt and Partridge, 2018). In support, empirical studies have found that genes with expression biased towards late life are less conserved than those highly expressed at young age across several tissues and mammalian species (Turan et al., 2019; Jia et al., 2018) Building on this, the antagonistic pleiotropy theory describes how genes with beneficial effects early in life can be maintained by selection even if they have pleiotropic negative effects later in life (Williams, 1957). In the disposable soma theory, the pleiotropic effect of more specific genes is described, that cause a trade-off between somatic maintenance and reproduction (Kirkwood, 1977), so that an increased, or early, investment in offspring is expected to come at the price of a shorter lifespan and vice versa (Kirkwood and Austad, 2000).

There are, however, exceptions to these expectations; possibly most notably within social insects, where reproductive castes exhibit relatively long lifespans compared to their sterile siblings (Keller and Genoud, 1997). This apparent lack of a trade-off between longevity and fecundity in social insects is at odds with expectations for the disposable soma theory. The longer life of queens compared to sterile castes might be explained by low extrinsic mortality due to the protection of a well-defended nest (Keller and Genoud, 1997; Negroni et al., 2016). The low extrinsic mortality of queens can in turn be expected to lead to a reduction of the selection shadow as more queens reach old-age, allowing efficient selection on genes that are important for somatic maintenance late in life.

In an attempt to understand the relationship between fecundity and longevity in social insects, several studies have investigated caste and age-specific expression of putative ageing genes in honeybees (Aamodt, 2009; Aurori et al., 2014; Corona et al., 2005, 2007; Seehuus et al., 2013), ants (Lucas et al., 2016; Lucas and Keller, 2018; Negroni et al., 2019; Von Wyschetzki et al., 2015) and termites (Kuhn et al., 2019; Elsner et al., 2018). One of these studies, which compared gene expression between young and old queens of the ant *Cardiocondyla obscurior*, identified several overlaps with ageing pathways known from *Drosophila melanogaster* (Von Wyschetzki et al., 2015). However, surprisingly, for many genes the ratio of expression level between old and young ant queens was reversed compared to *D. melanogaster*. Further studies comparing expression between castes and age-groups highlight the importance of several gene pathways for longevity in social insects that have previously been implicated in ageing, such as antioxidants (Aurori et al., 2014; Corona et al., 2005; Negroni et al., 2019; Kuhn et al., 2019), immunity (Negroni et al., 2019; Aurori et al., 2014; Lucas and Keller, 2018; Kuhn et al., 2019; Negroni et al., 2016), DNA and somatic repair (Kuhn et al., 2019; Aamodt, 2009; Lucas et al., 2016; Seehuus et al., 2013), respiration (Lockett et al., 2016; Corona et al., 2005), as well as the insulin/insulin-like growth factor (IGF) signaling (IIS) (Kuhn et al., 2019; Aurori et al., 2014) and the target of rapamycin (TOR) signalling pathways (Negroni et al., 2019; Kuhn et al., 2019). The IIS and TOR nutrient sensing pathways are of particular interest in this context, since their role in longevity and fecundity has been extensively studied in model organisms (Tatar et al., 2003; Partridge et al., 2011; Kenyon, 2010; Flatt and Partridge, 2018). These transcriptional studies offer insights into individual genes and their pathways that might be involved in ageing in social insects. However, a more holistic view of gene networks is likely to uncover further important genes as well as insights into transcriptional regulation. For example, a study of gene co-expression networks on mouse brains revealed that with age a decrease in the correlation of expression between genes occurred, showing that transcriptional dysregulation can lead to a significant reduction in gene connectivity (Southworth et al., 2009). These findings demonstrate the application of transcriptional studies for investigating whole pathways and gene networks and their wide-reaching implications for ageing. Furthermore, the extent at which a selection shadow may be reduced for old queens due to a reduction in extrinsic mortality has so far not been formally tested.

To address these questions we investigated transcriptomic data available for young and old queens of the polygynous ant, *C. obscurior* (Von Wyschetzki et al., 2015). These ant queens are relatively short-lived compared to most ant species (median lifespan: 16-26 weeks Kramer et al. 2015; Schrempf et al. 2005), which is in accordance with expectations for polygynous species, where extrinsic mortality is higher than in monogynous colonies (Keller and Genoud, 1997). Nevertheless, as for most ant species, *C. obscurior* queens (up to 48 weeks) outlive sterile workers that are expected to live around 12 to 16 weeks (Oettler and Schrempf, 2016). Importantly, consistently high reproductive output throughout their lives until immediately before death indicates no apparent reproductive senescence in these ant queens (Kramer et al., 2015). To test for signs of ageing in transcriptional regulation, we carried out a gene co-expression network analysis, in which we identified gene modules related to young mated (4 weeks) and old mated (18 weeks) queens and compared overall network connectivity. We also tested the hypothesis that, due to low extrinsic mortality, selection efficiency should not decline with age in queens. We found evidence for an array of anti-ageing mechanisms that are more tightly regulated in old queens. We could find no evidence for a selection shadow, indicating stable selection efficiency throughout an ant queen life.

## Results and Discussion

### Old-biased genes are not under weaker selection

Evolutionary theories of ageing predict weaker selection on genes which are expressed in old individuals due to low effective population size and reduced fecundity (Kirkwood and Austad, 2000; Flatt and Partridge, 2018). In ant queens, we may expect a reduction of this ‘selection shadow’ as low extrinsic mortality and lifelong, high fertility should lead to a stable effective population size up to old age. We tested this by estimating and comparing selection strength between three groups of genes. These were (i) old-biased genes n=46: significantly over-expressed in seven old (18 weeks) compared to seven young (4 weeks) *C. obscurior* queens; (ii) young-biased genes (n=96): significantly over-expressed in young compared to old queens; (iii) unbiased genes (n=2616): no significant difference in expression between young and old queens. To estimate direction and strength of selection, we measured dN/dS (ratio of nonsynonymous to synonymous substitution rates) for one-to-one orthologs with a set of 10 ant species (see methods). A dN/dS ratio *≈* 1 indicates neutral evolution, whereas values ≪ 1 signify purifying selection. We find no evidence for weaker purifying selection in old-aged queens, since dN/dS in old-biased genes (median: 0.084) is in fact significantly lower than in young-biased genes (median: 0.127; p-value = 0.016; Mann-Whitney U test; fig. 1), indicating increased purifying selection with age. Interestingly, dN/dS in young-biased genes is also significantly lower than in unbiased genes (median: 0.100; p-value = 2.2×10^*−*^4; Mann-Whitney U test), as has previously been reported for the ant, *Lasius niger* (Lucas et al., 2017). This is in contrast to published results for age-biased genes in humans, in which old-biased genes had a significantly higher dN/dS (median: 0.22) than young-biased (median: 0.09, p = 1.4×10^*−*50^), as would be expected for a reduction in purifying selection with age (Jia et al., 2018). This was confirmed by a further study on several mammalian tissues, in which an adjusted dN/dS metric correlated more strongly with expression in young compared to old individuals (Turan et al., 2019). To further test the ability of this method to detect a selection shadow in insects, we repeated the analysis for *D. melanogaster*. Age-biased gene expression was measured for a novel data set containing expression data for young (10 days) and old (38 days) female flies across two tissues (head and fat body) and different feeding regimes. Evolutionary rates were obtained for these genes from published analyses based on alignments of 12 *Drosophila* species (Consortium et al., 2007). In contrast to our results for ant queens but in agreement with expectations for a selection shadow, we find significantly higher dN/dS levels in old-biased fly genes (median: 0.060) compared to young-biased genes (median: 0.047; p=5.1×10^*−*^8; Mann-Whintey U test).

**Figure 1:**
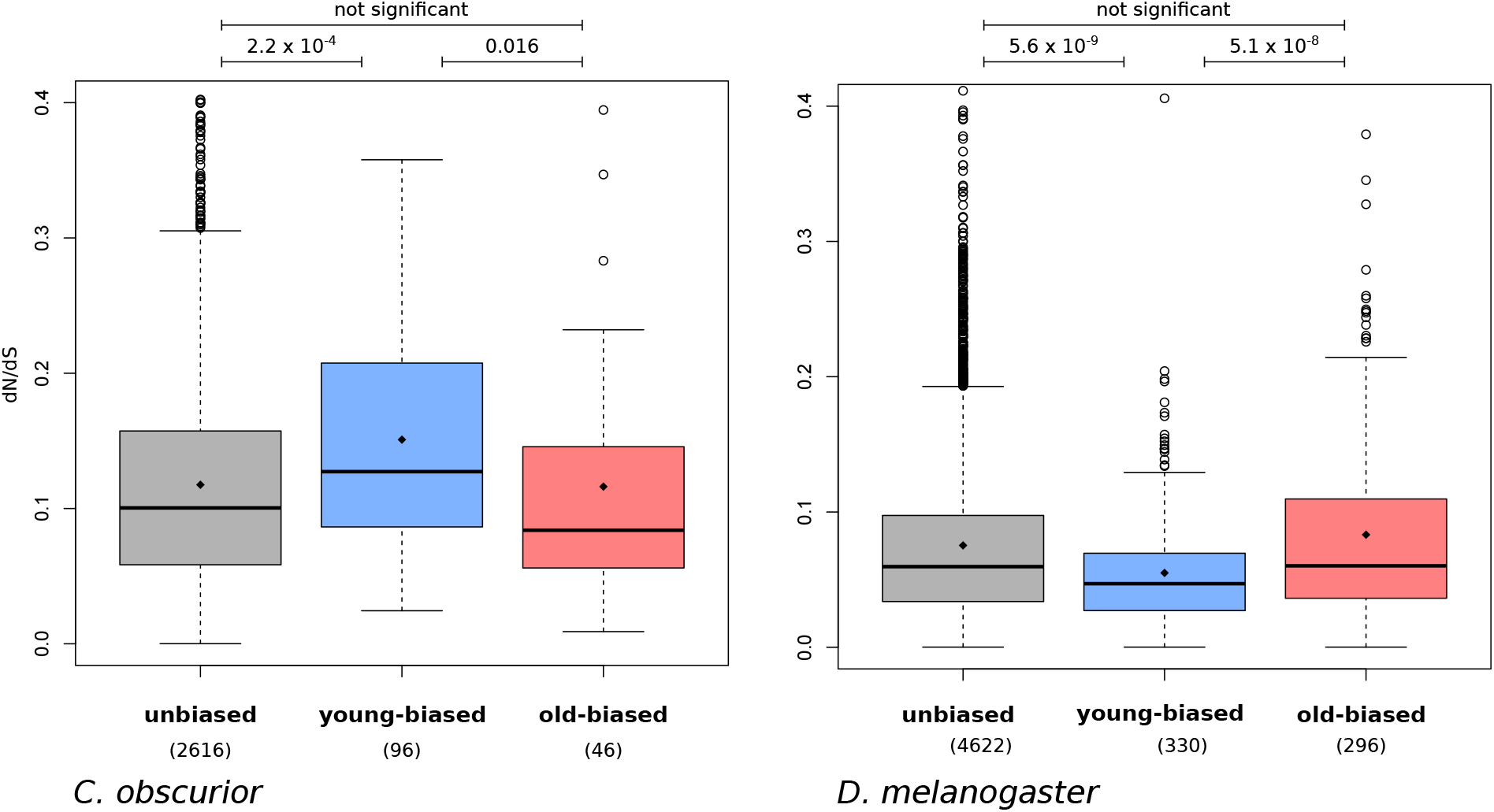
Evolutionary rates (dN/dS) in genes with unbiased expression, young-biased and old-biased expression in *C. obscurior* queens and *D. melanogaster* adult females. Significance was tested with Mann-Whitney U test.

We also investigated the numbers of ant genes that are under significant positive selection within old-biased compared to young-biased and unbiased genes, using a site test of the codeml suite (Yang, 1997). Contrary to expectations for weaker selection strength on old queens, we found no difference in the proportion of genes under positive selection between the three groups of genes (old-biased: 21.7%; young-biased: 21.9%; unbiased: 16.0%; Chi^2^ = 3.3; p = 0.19). The effect size of the observed difference in proportions of genes under positive selection between young- and old-biased genes is so low (cohen’s h: 0.003), that we assume the lack of significance is not due to a lack of power. The genes under significant positive selection in old-biased genes contain two regulatory genes (transcription factor and methyltransferase), an electron transport protein, a member of the COPI coatomer complex (important for protein transport) and *Notch* (Table 1). The latter is the central signalling protein within the Notch signalling pathway which is involved in tissue homeostasis and age-related diseases (Balistreri et al., 2016). Contrary to expectations based on evolutionary theories of ageing, these results suggest selection is not weaker on genes expressed mainly in old queens. We speculate that high fertility in old queens, coupled with an overall low extrinsic mortality, which is typical for social insects (Negroni et al., 2016; Keller and Genoud, 1997), may reduce the selection shadow in *C. obscurior* queens, leading to similar selection strength throughout their fertile life.

**Table 1:** Old-biased genes under significant positive selection.

| Gene | Ortholog | Putative Function |
| --- | --- | --- |
| Cobs_01221 | uncharacterised | unknown |
| Cobs_04278 | FBgn0002121 (l(2)gl) | polarity of neuroblasts and oocytes |
| Cobs_06663 | FBgn0085424 (nub) | transcription factor |
| Cobs_08231 | FBgn0004647 (Notch) | tissue homeostasis |
| Cobs_08620 | FBgn0027607 (Dymeclin) | organisation of Golgi apparatus |
| Cobs_09212 | FBgn0033686 (Hen1) | methyltransferase, methylates siRNA & piRNA |
| Cobs_11651 | FBgn0036714 (CG7692) | unknown function |
| Cobs_12452 | FBgn0008635 (βCOP) | subunit of the COPI coatomer complex, transport from Golgi to ER |
| Cobs_16420 | FBgn0034745 (CG4329) | unknown |
| Cobs_16765 | Cytochrome b561 domain-containing protein 1 (Q8N8Q1) | electron transport protein |

### Increased connectivity in old ant queens

In old queens, we expected to find little evidence for age-related transcriptional dysregulation in the form of reduced correlation of gene expression, as previously reported for ageing mouse brains (Southworth et al., 2009). We investigated this by measuring gene connectivity separately within old queens and within young queens, using the *softConnectivity* function of the WGCNA package (Langfelder and Horvath, 2008). This connectivity describes the total strength of correlations that a gene possesses with all other genes in a gene co-expression network (GCN; Langfelder and Horvath 2008) and is thought to correlate positively with gene essentiality (Carlson et al., 2006). In fact, we find gene expression connectivity to be significantly higher in older queens (median: 145.3) than within young queens (median: 142.5; effect size: 0.255; p = 4.3×10^*−*29^; Wilcoxon signed-rank test), suggesting an increased regulation of gene networks in older queens.

The more highly connected genes in older queens (1471 genes with connectivity fold change > 2) are enriched for GO term functions (FDR < 0.1) related to protein synthesis, transcription, purine synthesis, cellular respiration and ATP metabolism (Table S1). Most of the 20 genes with the strongest increase in connectivity in old queens (4.8-7.1 fold increase) compared to young queens are involved in transcriptional regulation (7 genes) or protein homeostasis (6 genes; Table S2). For example, a member of the 26S proteasome complex, important for the degradation of misfolded proteins, is the gene with the highest increase in connectivity in old queens. As has been shown for several organisms, including humans (Lee et al., 2010), yeast (Kruegel et al., 2011) and *C. elegans* (Vilchez et al., 2012), increased proteosome activity can extend lifespan by reducing proteotoxic stress (López-Otín et al., 2013). An increase in connectivity of fatty-acid synthase 3 may have implications for colony communication. Further highly connected genes include ribosomal proteins or genes involved in the correct folding, post-translation modification or transport of proteins. The genes with highly increased connectivity in old ant queens, which are involved in transcriptional regulation, include two transcription factors, a transcritional coregulator (*taranis*), and four mRNA regulators. These results suggest that, contrary to expectations for ageing individuals, increased transcriptional regulation and protein homeostasis takes place in old queens.

### Co-expression modules related to age

We constructed a signed, weighted gene co-expression network (GCN, Langfelder and Horvath 2008) based on the correlation of normalised gene expression across all 14 samples (7 young queens & 7 old queens). Within the GCN, genes could be grouped into 27 modules, within which gene expression was especially strongly correlated (Fig. 2). To determine the importance of these modules for old and young queens, we first calculated eigengene expression based on the first principal component of each module. We then correlated eigengene expression of each module with the binary trait ’age’ (young & old). Five of the modules were significantly, positively correlated with age (p < 0.05; FDR < 0.1), indicating an overall higher expression of these modules in old compared to young queens. Three modules were significantly, negatively correlated with young queens, indicating a downregulation in old queens. To validate these correlations, we analysed difference in expression of genes between old and young queens (log_2_[expression_*old*_/expression_*young*_]) within each of these modules. Accordingly, the median log_2_-fold-change in expression was greater than zero in each of the old-biased modules (0.148 to 0.340) and less than zero within the young-biased modules (−0.376 to -0.249; Fig. S1). Four of the five old-biased modules (1, 2, 3 and 5) belonged to a larger cluster within the GCN, which is quite distant from the cluster containing the young-biased modules (6, 7, 8; Fig. 2). module 4 (old-biased), on the other hand, forms a more distinct cluster, adjacent to the young-biased cluster. The old-biased modules contained several genes that had previously been identified as upregulated in old queens via standard differential expression analysis (Von Wyschetzki et al., 2015) but contained no genes that were upregulated in young queens. The opposite was true for young-biased modules, thus confirming the validity and compatibility of both methods (Fig. 2(b)).

**Figure 2:**
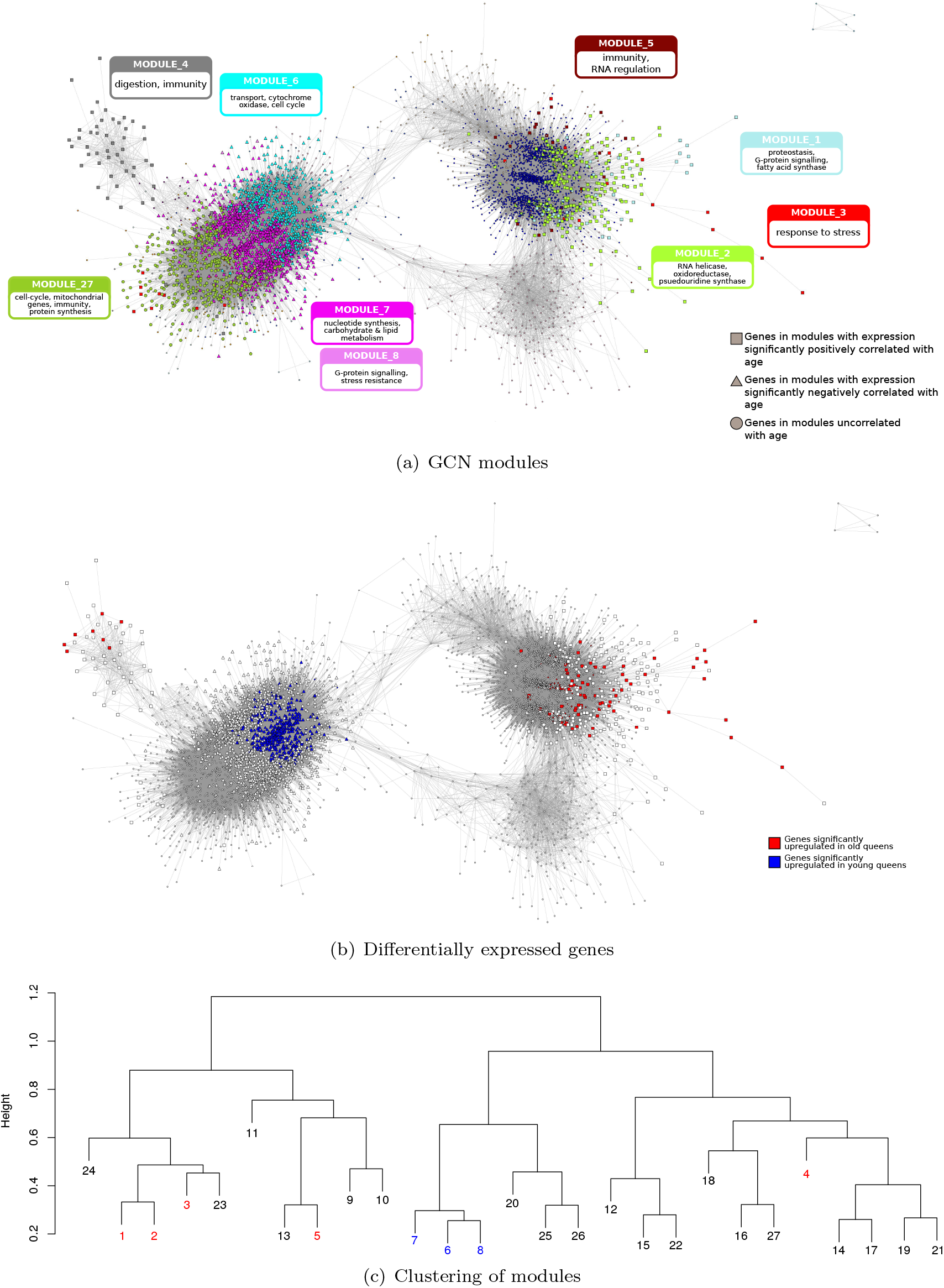
Gene co-expression network (GCN). (a & b) Graphical representation of the gene co-expression network, containing only the most strongly connected genes (n = 5442). In (a) genes are coloured according to the modules to which they belong. The main enriched functions (based on hubs and GO terms) of the 9 discussed modules are labelled (see text for more details). In (b) genes are coloured according to their differential expression; red: over-expressed in old queens; blue: over-expressed in young queens; white: not differentially expressed. In both representations, genes in modules significantly related to old queen expression are depicted as squares, and those significantly related to young queens are triangles; all other genes are represented by circles.(c) Clustering dendogram of modules; height represents dissimilarity based on topological overlap. Modules significantly related to age are highlighted in red (positive correlation) and blue (negative correlation). Higher resolution image available in the online version.

However, importantly, the GCN analysis also allowed the identification of many additional age-related genes that can not be identified by standard differential expression analyses. For example, module 1, which has the strongest association with old queens (*ρ* = 0.96; p = 5.3×10^*−*8^; FDR = 1.4×10^*−*6^; Pearson correlation), contains 109 genes, of which only 41 are individually significantly differentially expressed between old and young queens. Similarly, module 6, which is strongly negatively associated with old queens (*ρ*r = -0.90; p = 9.4×10^*−*6^; FDR = 1.3×10^*−*4^; Pearson correlation), contains 970 genes, of which 240 were identified as individually significantly upregulated in young queens (Von Wyschetzki et al., 2015). In the following section, we describe these eight age-biased modules in terms of their functional enrichment and detail the top hub genes (genes with the highest intramodular connectivity) within these modules.

#### Old-biased modules

The most highly connected hub genes in ***module 1***, the module most strongly upregulated with age (*ρ* = 0.96; p = 5.3x^10*−*8^; FDR = 1.4×10^*−*6^; 109 genes; Fig. 3), include three genes with functions related to maintaining and restoring proteostasis in old queens (Table S3), the loss of which has been described as one of the hallmarks of ageing (López-Otín et al., 2013). These are: a member of the TRAPP complex, important for protein transport, *Socs44A*, a gene involved in ubiquitination and *GRXCR1*, responsible for the post-transcriptional S-glutathionylation of proteins, a modification which is often triggered as a defence against oxidative stress (Dalle-Donne et al., 2009). The top hubs of this module also include two genes which encode integral members of the G-protein signalling pathway, namely, a Rho guanine nucleotide exchange factor and a G-protein *α*-subunit. The most connected gene within this hub is a fatty-acid synthase which may play an important role in colony communication. This module is enriched for a GO term related to the regulation of transcription (Table S4).

**Figure 3:**
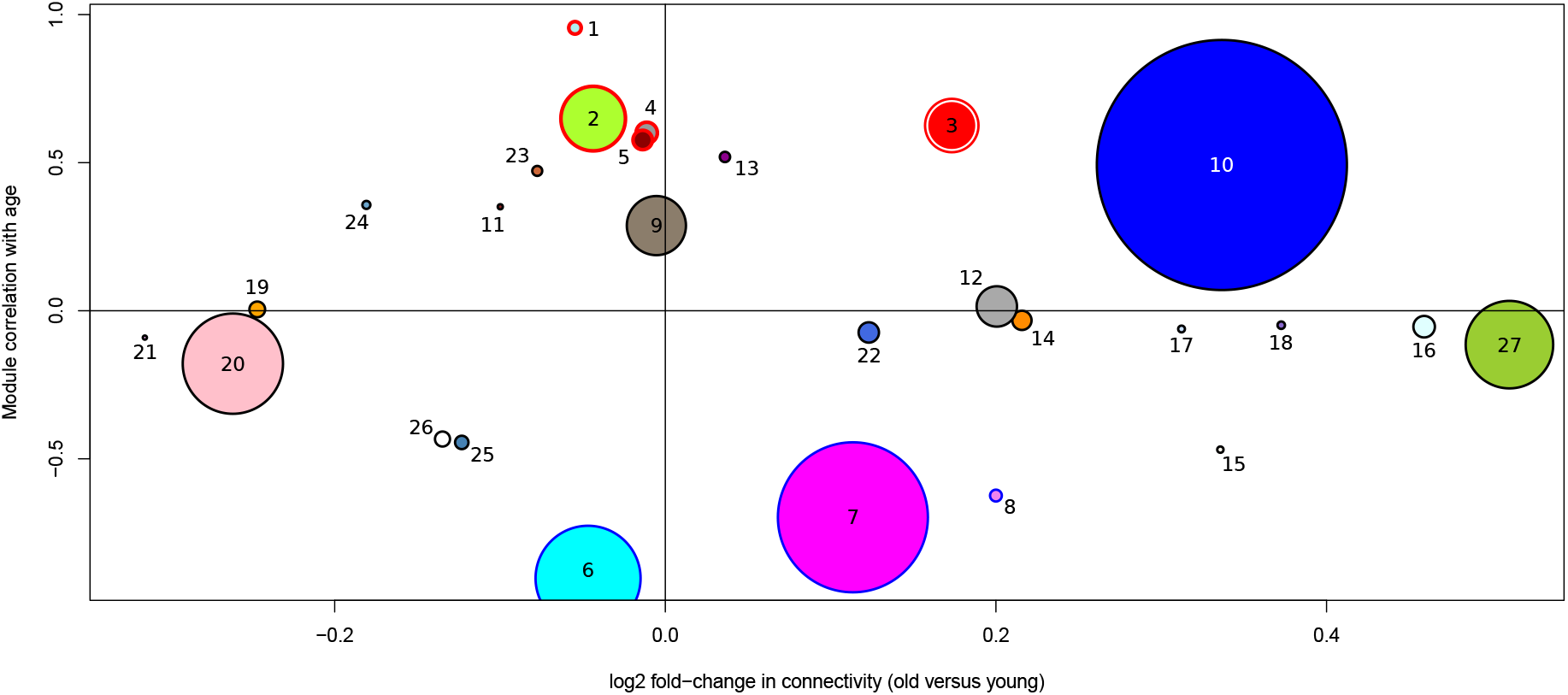
Correlation of GCN modules with age and their change in connectivity between old and young queens. A positive correlation with age (y-axis) signifies an upregulation of a module in old queens. A positive log2foldchange in connectivity (x-axis) represents a higher connectivity in old queens. Modules are labelled with their assigned module numbers. Sizes of dots represent relative number of genes within modules. Modules with red outlines are significantly upregulated and modules with blue outlines are significantly downregulated in old queens compared to young queens.

***Module 2*** (596 genes;upregulated with age: *ρ* = 0.65; p = 0.012; FDR = 0.080) contains hub genes coding for proteins with diverse functions, including an RNA helicase, a maternal protein, a protein with oxidoreductase activity and a pseudouridine synthase (Table S3).

***Module 3*** (433 genes; upregulated with age; *ρ* = 0.63; p = 0.017; FDR = 0.080) is particularly interesting since it is not only upregulated with age but, on average, gene members of the module are more strongly connected within old than in young queens (Fig. 3). Hub genes indicate this module is important for responses to age-related stress, especially processes related to a maintenance of proteostasis (Table S3). For instance, the top 10 hubs contain the endoplasmic reticulum (ER) stress protein, disulfide-isomerase, which reacts to protein misfolding and oxidative stress (Laurindo et al., 2012), as well as *fringe*, which modulates Notch signalling, a pathway important for regulating tissue homeostasis and implicated in ageing related diseases (Balistreri et al., 2016). A further hub is a trehalose transporter, orthologous to *tret1-2*, indicating that the transport of trehalose (the main haemolymph sugar in insects) from fat body to other tissues is well regulated in old queens (Kanamori et al., 2010). This may have a positive effect on survival, since trehalose treatment increases longevity in *C. elegans* (Honda et al., 2010).

The top 10 hub genes in ***module 4*** (186 genes; *ρ* = 0.61; p = 0.021; FDR = 0.080) fulfil various functions, such as the digestive enzymes alpha glucosidase and chymotrypsin-1, indicating a possible modification in diet with age (Table S3). The third most connected gene within this module is orthologous to *pirk* in *D. melanogaster* (involved in the negative regulation of the immune response; Kleino et al. 2008), indicating the immune system may be downregulated with age in *C. obscurior*. Interestingly, long-lived flies also tend to downregulate the induction of immune effector genes (Fabian et al., 2018; Loch et al., 2017). This module is enriched for the GO term transmembrane transport (Table S4).

***Module 5*** (169 genes; *ρ* = 0.58; p = 0.028; FDR = 0.095) may be important for controlling the immune system since two hub genes (Table S3), the COMM domain containing protein 8 (COMMD8) and the WD40 domain containing angio-associated migratory cell protein (AAMP), are both known to inhibit the transcription factor NF-*κ*-B (Burstein et al., 2005; Bielig et al., 2009). An upregulation of NF-*κ*-B occurs with ageing and its inhibition, as apparently occurs within this module, can reduce the effects of senescence (Tilstra et al., 2012). Interestingly, COMMD8 is also characterised by a strong increase in connectivity (1.68 fold change), indicating its heightened importance in old queens. Further functions of this module may be related to RNA regulation, as evidenced by the hub gene *eyes absent*, a transcription factor with importance in embryonal eye development in *D. melanogaster* (Bonini et al., 1998). Based on the ten nearest neighbours in the *C. obscurior* GCN, *eyes absent* may regulate several enzymes involved in post-transcriptional processes, such as mRNA export from the nucleus (*sbr*, Cobs 03187), and tRNA modification (*Tgt* : Cobs 16650; *HisRS* : Cobs 01013; CG3808: Cobs 18201).

#### Modules downregulated with age

***Module 6*** (970 genes) is the module most strongly down-regulated with age (*ρ* = -0.9; p = 9.4×10^*−*6^; FDR = 1.3×10^*−*4^) and is enriched for the GO terms transmembrane transport and potassium ion transport (Table S4). Interestingly, the top 10 hubs contain three genes with no detectable homology to any protein in the uniprot arthropod database (Table S3). Otherwise, the functions of hub genes in this module span various functions, such as cell-cell interactions, cytochrome oxidase, an odorant receptor and a negative regulator of the cell cycle.

***Module 7*** (1385 genes; *ρ* = -0.7; p = 0.006; FDR = 0.050) has several enriched functions in the nucleotide synthetic process, oxidoreductase activity, carbohydrate and lipid metabolism, ATP metabolic processes, cofactor and coenzyme binding (Table S4). Accordingly the top hubs in this module contain a thioredoxin, a proteasome subunit (*α*6) and two genes involved in ubiquitination (*STUB1* and *Ubc6* ; Table S3).

***Module 8*** (103 genes; *ρ* = -0.62; p = 0.018; FDR = 0.080) is enriched forthe function G-protein coupled receptor activity (Table S4). The top hub gene in this module (intraconectivity 0.90), Cobs 08138, is orthologous to the

(Friedrich and Jones, 2016). Interestingly, mutant flies, carrying P-element insertions in one of these *methuselah* genes, live 35% longer and are significantly more resistant to stresses than wild-types (Lin et al., 1998). There are indications that these effects on lifespan and stress response may represent the ancestral function of methuselah receptors in *Drosophila* (Araújo et al., 2013). A similar function of the *methuselah* ortholog in *C. obscurior* would explain how a reduction in expression within older queens may facilitate life extension and greater stress resistance.

We also examined ***module 27*** (808 genes) in more detail since it shows the strongest increase in connectivity in old compared to young queens (1.47 fold) of all modules (Fig. 3), suggesting an increased regulation of this module with age. The functions connected to this module, based on hubs (Table S3), increases in connectivity (Table S5) and GO terms (Table S4), indicate that in old queens an increased regulation of cell-cycle, mitochondrial genes, immunity genes, transcriptional genes and members of the protein synthesis machinery takes place, which is in stark contrast to the expected gene expression hallmarks of ageing in multicellular eukaryotes (Frenk and Houseley, 2018).

#### Robustness of GCN

Since our sample size of 14 is one lower than the recommended minimum of 15, we confirmed the robustness of our results by adding further samples from the same study (Von Wyschetzki et al., 2015). For this, we incorporated expression data from 7 old queen samples that had mated with sterile males (‘sham-mated’) and then created 8 further GCNs, 7 of which contained one sham-mated queen (total n = 15) and one GCN containing all 7 sham-mated queens (n = 21). We used preservation statistics (Langfelder et al., 2011) to compare the modules of our GCN with these larger GCNs. Within each module, correlation, adjacency, connectivity and variance explained by the eigennode are compared between all nodes, and for each statistic a z-score is calculated based on 200 permutations. A composite z-summary of these statistics is calculated, whereby a threshold of 2 is deemed as necessary for classing a module as preserved, while a score greater than 10 offers strong evidence for module preservation. In each comparison against the 8 additional, larger GCNs, our age-biased modules scored at least 10, offering strong support that our GCN is not affected by a limited sample size.

### Old-biased module hubs are highly conserved

We investigated evolutionary rates of the most connected genes within the old- and young-biased modules. Hub genes (intraconnectivity > 50%) of the five old-biased modules have significantly lower rates of protein evolution (dN/dS median: 0.081) than hubs in young-biased modules (median: 0.118; p = 6.0×10^*−*4^) or compared to all lowly connected genes (intraconnectivity < 50%; median: 0.101; p = 0.017; Fig. 4). We investigated the influence of expression levels on these results, since highly expressed genes are often found to be under stronger purifying selection (Drummond et al., 2005). However, expression levels, based on mean normalised read counts among all 14 samples, do not differ between hub genes of old-biased (mean: 291.5) and young-baised genes (mean: 326.8; W = 3160, p-value = 0.18). These results suggest the hub genes of old-biased modules are highly constrained by strong purifying selection.

**Figure 4:**
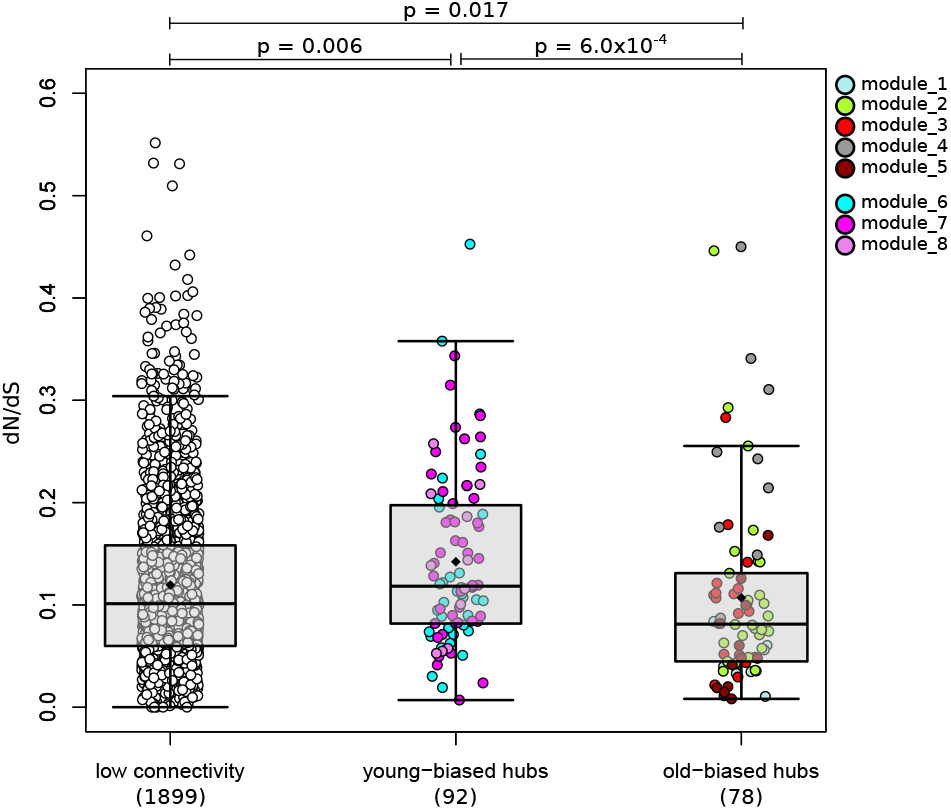
dN/dS rates in hub genes within young- and old-biased modules compared to lowly connected genes. Each dot represents a gene, which are coloured by the module membership. Whiskers of the boxplots represent up to 1.5 times the interquartile range. Black diamonds are means, and horizontal bars within the boxes are medians. Hub genes have an intraconnectivity > 50%; lowly connected: < 50%.

### Conclusions

Evolutionary theory of ageing predicts a selection shadow on genes expressed late in life due to a reduction in effective population size with increasing age caused by extrinsic mortality (Kirkwood and Austad, 2000). We expected to find a reduced selection shadow in *C. obscurior* queens, as ant queens generally experience low extrinsic mortality. In support, we find compelling evidence for strong purifying selection on old-biased genes (significantly upregulated in 7 old compared to 7 young queens), for which evolutionary rates (dN/dS) are significantly lower than young-biased genes. In contrast, we find evidence of a selection shadow in *D. melanogaster* where dN/dS is significantly higher for old-biased genes. Our results suggest, therefore, that *C. obscurior* queens are not affected by a selection shadow, so that genes important at old age can not be expected to accumulate deleterious mutations at an increased rate compared to early-acting genes. This offers an explanation for the apparent lack of ageing and the high reproductive output of old ant queens.

Furthermore, we were interested in understanding whether *C. obscurior* queens show signs of ageing, especially within transcriptional regulation. This is a particularly intriguing question since the reproductive fitness of these ant queens remains high until old age, although they outlive their sterile siblings (Oettler and Schrempf, 2016). In fact, our analysis of co-expression networks in *C. obscurior* queens uncovers a significant increase in gene connectivity in old queens. This result offers evidence for an increased transcriptional regulation, especially in genes that are themselves involved in transcriptional regulation, as well as several genes involved in protein synthesis and degradation, which are important mechanisms for counteracting symptoms of ageing (Frenk and Houseley, 2018). Also, the analysis of old-biased modules (clusters of highly correlated genes, upregulated with age) within the GCN revealed an increase in expression and connectivity of genes involved in proteostasis, stress response, and transcriptional regulation (Fig. 2(a)), offering further support for well-regulated anti-ageing mechanisms. The hub genes within these old-biased modules are more highly conserved than hubs of young-biased modules, indicating strong purifying selection acting on these important central regulators.

In summary, we find no evidence of ageing in transcriptional regulation in *C. obscurior* queens. Low extrinsic mortality may allow selection to shape genes important at old age, which is evident in low divergence rates (dN/dS) of the hubs of old-biased modules. Well regulated molecular mechanisms likely allow the ant queens to counteract any symptoms of ageing, thus maintaining high reproductive fitness throughout life. We suggest further transcriptional studies into the short period directly before death when the reproductive output of *C. obscurior* queens decreases (Heinze and Schrempf, 2012; Kramer et al., 2015), which we expect to illuminate processes of transcriptional ageing. Transcriptional studies of other ant species are necessary to investigate the generality of our findings. In monogynous ants, for example, in which individual queens are less dispensible, we would expect to observe an even weaker selection shadow. Also, *C. obscurior* queens are relatively short-lived compared to other ant species. Selection strength on age-biased genes of extremely long-lived queens may be less affected by reductions in effective population sizes due to longer generation times. Further detailed research on individual pathways is important to understand how an upregulation of anti-ageing mechanisms occurs; especially proteomic analyses may reveal the true relationships between pathway members.

## Methods

### Data set

Genome and proteome sequences of the *C. obscurior* genome, version 1.4, were obtained from the hymenopteragenome.org website (accessed July 2018; Elsik et al. 2015). We estimated gene functions based on orthology, primarily to *D. melanogaster*, as well as PFAM domains and GO terms. Putative protein functions were based on descriptions in the flybase (Thurmond et al., 2018) and UniProt (Consortium, 2018) databases, unless otherwise stated. We calculated orthology to *D. melanogaster* with the method of reciprocal best blast hit (Rivera et al., 1998). For this, the proteomes of *C. obscurior* and *D. melanogaster* (v. 6.21; obtained from ftp://ftp.flybase.net/releases/current/dmel_r6.21/fasta/; accessed June 2018) were blasted against each other using blastp (BLAST 2.7.1+; Camacho et al. 2009) and an e-value threshold of 1e^*−*5^. Reciprocal best blast hits were extracted from the output files using a custom perl script. Where no orthology could be detected using this method, protein sequences were blasted against the swissprot database with blastp (version 2.7.1+; Altschul et al. 1990) and the best hit was retained with an evalue < 0.05. Protein sequences were annotated with PFAM domains using pfamscan (Mistry et al., 2007), to which GO terms were mapped with pfam2GO (Mitchell et al., 2014). Published RNAseq data were obtained for 7 old (18 weeks) and 7 young (4 weeks) ant queens from NCBI (Von Wyschetzki et al., 2015). These queens had each been individually reared from pupal stage in separate experimental colonies, each containing 20 workers and 10 larvae, originating from the genome reference population in Bahia, Brazil (Von Wyschetzki et al., 2015; Schrader et al., 2014). Fastq files were mapped to the *C. obscurior* genome (version 1.4) with hisat2 (Kim et al., 2019) using default parameters. We then indexed and sorted sam files using samtools (version 1.7, Li et al. 2009) and generated counts per gene using htseq-count (Anders et al., 2015). All statistical analyses on these counts were carried out in R (version 3.5.1, R Core Team 2018). Where necessary, we corrected for multiple testing with the p.adjust function, using the fdr method (Benjamini and Hochberg, 1995). A total of 10 339 genes were expressed in at least two individuals with a read count of at least 10. This subset of genes was used for all analyses.

### Determining age-biased expression

Within this subset of 10 339 genes, we identified genes with age-biased expression by comparing expression in the 7 old to the 7 young samples. This was carried out with the R package DESeq2 at default settings (Love et al., 2014). Genes with an adjusted p-value < 0.05 were deemed either old- or young-biased. All other genes were classified as unbiased.

### Molecular evolution and selection analyses

In order to carry out evolutionary analyses, we first determined orthology between the proteomes of *C. obscurior* and 9 further ant species, which we either downloaded from the hymenopteragenome.org website (accessed August 2020; Elsik et al. 2015): *Atta cephalotes, Pogonomyrmex barbatus, Solonopsis invicta* and *Wasmannia auropunctata*; or NCBI (accessed August 2020): *Monomorium pharaonis, Temnothorax cuvispinosus, Temnothorax longispinosus, Vollenhovia emery*. Data for *Crematogaster levior* were obtained from the authors of the genome publication upon request (Hartke et al., 2019). Orthology was determined with OrthoFinder (Emms and Kelly, 2015) at default settings. We chose orthologous groups that contained single gene copies within each of the 10 species. Protein sequences of each ortholog set were aligned with prank (version 170427, Löytynoja 2014) at default settings. The corresponding CDS sequences were aligned using pal2nal (Suyama et al., 2006). CDS alignments were trimmed for poorly aligned codon positions with Gblocks (version 0.91b) with the following parameters: -t=c -b2=6 -b3=100000 -b4=1 -b5=h. We calculated dN/dS ratios using the null model of codeml in the PAML suite (Yang, 1997), using the following tree based on a published ant phylogeny (Ward et al., 2015): (((((((Tlon, Tcur),Clev),Veme),Cobs),(Waur, Acep)),(Mpha, Sinv)),Pbar) dN/dS ratios were used for analyses only if dS < 3. dN/dS ratios were compared between old-biased, young-biased and unbiased genes using the Mann-Whitney test with the R function *wilcox*.*test*.

In order to detect genes that contain codon sites under positive selection, we performed a likelihood-ratio test (LRT) between models 7 (null hypothesis; dN/dS limited between 0 and 1) and 8 (alternative hypothesis; additional parameter allows dN/dS > 1) of the codeml program within the PAML suite (Yang, 1997). For this we used runmode 0, model 0 and set ’NSsites’ to 7 & 8.

### Gene co-expression analysis

The expression counts data were normalised using the built-in median of ratios method implemented by default in DESeq2 (version 1.22.2, Love et al. 2014) and then transposed to a matrix containing genes in columns and samples in rows. With the reduced set of 10 339 genes, we created a signed weighted gene co-expression network using the WGCNA package (version 1.68, Langfelder and Horvath 2008) that incorporated expression values from all 14 queen samples (7 young and 7 old). We followed the standard stepwise protocol (https://horvath.genetics.ucla.edu/html/CoexpressionNetwork/Rpackages/WGCNA/Tutorials/), using a soft power of 14 and the biweight midcorrelation function for calculating coexpression similarity. Minimum module size was set at 30 and resulting modules with a correlation of at least 0.75 were merged. Hub genes within modules were determined based on the intra-modular connectivity, which we calculated with the `intramodularConnectivity` function on the adjacency matrix, that was produced during the WGCNA pipeline. Age-biased modules were identified by correlating (pearson) the eigengene of each module with the binary trait young/old. FDRs were calculated with the p.adjust function, and modules with an FDR < 0.1 were considered significantly related to age.

To compare connectivity between young and old queens, we calculated connectivity with the `softConnectivity` function separately within the young and the old queen expression data. We used the same soft power value of 14 and the biweight midcorrelation function.

To create a visualisation of the GCN, the topological overlap matrix was reduced to only contain genes with a topological overlap of at least 0.1 to at least one other gene. Edge and node files were created with the WGCNA function `exportNetworkToCytoscape`, using a threshold of 0.1. All further visualisations of the network were conducted in Cytoscape (v. 3.7.2, Shannon et al. 2003).

To test the robustness of our GCN, we created 7 additional GCNs each with one extra sample taken from the sham-mated queens previously published within the same data set as our main data used here (Von Wyschetzki et al., 2015). We also created one larger GCN containing all 7 sham-mated queens, therefore containing 21 samples. Each additional GCN was created with the same parameters as our original GCN and then compared to our original GCN with the built-in WGCNA-function, *modulePreservation* and the *Zsummary* statistic was calculated. This composite z-score combines several comparative statistics, such as adjacency, connectivity and proportion of variance explained, with a score of 10 suggested as a threshold for strong evidence of module preservation (Langfelder et al., 2011).

### GO enrichment

GO term enrichment analyses were carried out with topGO (version 2.34.0; Alexa and Rahnenfuhrer 2018) on the “biological process” category, using the classic algorithm. Node size was set to 5, Fisher statistic was implemented and we only kept GO terms that matched at least 3 genes and with a p-value < 0.05. An FDR was added using the R function p.adjust and the method “fdr” (Benjamini and Hochberg, 1995); GO terms with an FDR < 0.1 were described in the text.

### D. melanogaster data set

To estimate evidence of a selection shadow in

*D. melanogaster*, we accessed a recently compiled, but so far unpublished, RNAseq data set (SRA accession: PRJNA615318). This data set comprised RNAseq of 34 samples of 5 pooled flies. We used y^1^,w^1118^ mutant flies (full genotype: yw; +/+; +/+). These flies were maintained in laboratory conditions at 25°C, 12h:12h light:dark and 60% relative humidity.

#### Experimental setup

Adult virgin females and males were collected separately, and 3 days later they were pooled together to freely mate. Eggs were laid in a controlled density (50-100 eggs per bottle) and developed until the adult stage in the same conditions as mentioned above. After eclosion, the offspring adult flies matured for one day. On the second day after eclosion, female and male flies were collected and transfered to a demographic cage. Each cage contained 130 females and 70 males. Once cages were set up, they were divided into four groups, which consisted of 4 different diet treatments. The diet treatments differed only in the content of yeast (20, 40, 80 or 120g) present in the fly food; the other ingredients were added in the same quantities in all diets (1L water, 7g agar, 50g sugar, 10mL 20% nipagin and 6mL propionic acid). All cages were maintained in the same conditions as described above.

### Sampling and RNA extractions

Female flies were sampled at two time points: 10 days (young) and 38 days (old). For each time point, sampling and dissections were done between 1 pm and 6 pm. Two groups of 5 females each (2 replicates) were anesthetized in the fridge (approximately 4°C), and afterwards fat bodies were dissected in ice-cold 1x PBS. To guarantee that we sampled the entire fat body, we decided to use in this experiment fat bodies still attached to the cuticle – usually referred to as fat body enriched samples – because the cuticle is transcriptionally inactive. In ice-cold PBS, the female fly abdomens were opened, and the organs were carefully removed. Once the fat body tissue was clean, the abdomen cuticle was separated from the thorax. The fat body enriched tissues were transferred into Eppendorf with 200µL of homogenization buffer from the RNA isolation kit (MagMAX™-96 Total RNA Isolation Kit from Thermo Fisher). The tissues were homogenized and stored at -80°C until RNA extraction. To sample head transcriptomes, flies were transferred to Eppendorfs and snap-frozen with liquid nitrogen. Then the Eppendorfs were vigorously shaken to separate the heads from the bodies. The heads were then transferred into an Eppendorf containing 200µL of homogenization buffer, from the RNA isolation kit. As described above, tissues were homogenized in the solution and kept at -80°C until RNA extraction. All extractions were done using the MagMax robot from Thermo Fisher and the MagMAX™-96 Total RNA Isolation Kit. In this experiment there is a total of 34 samples: 2 time points X 4 diet treatments x 2 tissues = 16 groups, for each group we have 2-3 replicates (all groups have 2 replicates except for the second time point for 2% yeast diet, where we have 3 replicates). The sequencing of the RNA samples was done in BGI, Hong Kong, China. The samples were sequenced (paired-end, 100bp) on an Illumina HiSeq 4000 platform. Gene counts were generated in the same manner as for *C. obscurior* using genome version 6.21 (obtained from ftp://ftp.flybase.net/releases/current/dmel_r6.21/fasta/; accessed June 2018).

## Acknowledgements

This paper was written as part of the research carried out by the DFG Collaborative Research Unit (RU) ‘Sociality and the Reversal of the Fecundity-longevity Trade-off’ (DFG FOR2281, www.so-long.org), and we thank the members of the RU for stimulating discussions. MCH is supported by a DFG grant BO2544/11-1 to EBB. JO and LMJN are supported by DFG grant OE549. MR and TF were supported by the Swiss National Science Foundation (SNSF) (grants 310030E-164207 and 31003A 182262 to TF) and the Novartis Foundation for Medical-Biological Research (grant 19B149 to TF).

## Author information

### Affiliations

Institute for Evolution and Biodiversity, University of Münster, Münster, Germany: Erich Bornberg-Bauer & Mark C Harrison

Department of Protein Evolution, Max Planck Institute for Developmental Biology, Tübingen: Erich Bornberg-Bauer

Institute for Zoologie/Evolutionary biology, University of Regensburg, Regensburg, Germany: Luisa M. Jaimes-Nino & Jan Oettler

Department of Biology, University of Fribourg, Fribourg, Switzerland: Marisa Almeida Rodrigues & Thomas Flatt

### Contributions

MCH, EBB conceived and initiated the project. MCH, EBB and JO designed the study. MCH wrote the manuscript and carried out most analyses. JR assisted with dN/dS analyses. MCH and JO interpreted ant data, all authors interpreted comparative data. LMJ assisted in the interpretation of GO term enrichment analyses. TF and MAR generated fly data and helped analyse them. MCH wrote the manuscript which was revised and approved by all authors.

### Corresponding authors

Correspondence to Erich Bornberg-Bauer and Jan Oettler.

### Data Availability

Ant queen data are already published (Von Wyschetzki et al., 2015) and available at SRA under accessions: PRJNA293450 & PRJNA284224. *Drosophila* data are deposited on SRA under accession: PRJNA615318. Scripts are available on the github: https://github.com/MCH74/AgeingInCardiocondyla

## Supplementary figures

**Figure S1:**
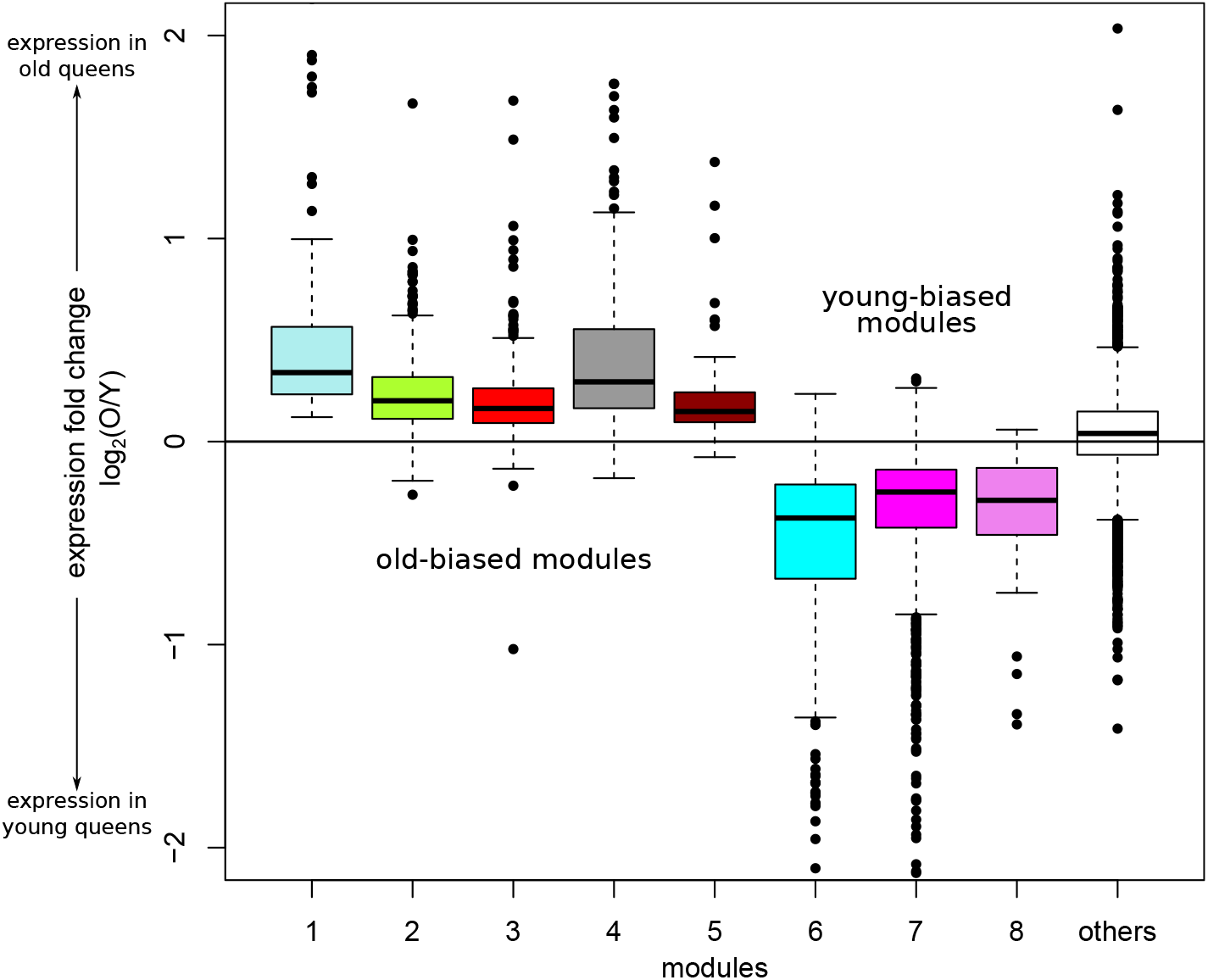
Fold change of expression in old compared to young queens in each of the significant modules (log_2_(old/young)). Correspondingly, genes within the ‘old-biased’ modules (1-5) show log_2_-fold-change of expression > 0 (medians: 0.340, 0.201, 0.163, 0.294, 0.148, respectively) and ‘young-biased’ modules (6-8) contain genes with negative log_2_-fold-change of expression (medians: -0.376, -0.249, -0.289, respectively). Expression fold change for genes of all other modules (white plot, right-most), on the other hand, has a median close to zero (0.040).

## Supplementary Tables

**Table S1:** GO terms (Biological Process) significantly enriched within genes with a connectivity fold change greater than 2 in old compared to young queens.

| GO.ID | Term | Annotated | Significant | Expected | pvalue | FDR |
| --- | --- | --- | --- | --- | --- | --- |
| GO:0006518 | peptide metabolic process | 168 | 67 | 20.85 | 1.1e-20 | 3.4e-18 |
| GO:0043043 | peptide biosynthetic process | 163 | 65 | 20.23 | 4.5e-20 | 6.4e-18 |
| GO:0006412 | translation | 160 | 64 | 19.85 | 7.5e-20 | 7.7e-18 |
| GO:0043604 | amide biosynthetic process | 166 | 65 | 20.6 | 1.4e-19 | 9.8e-18 |
| GO:0043603 | cellular amide metabolic process | 175 | 67 | 21.71 | 1.6e-19 | 9.8e-18 |
| GO:1901566 | organonitrogen compound biosynthetic pro... | 270 | 83 | 33.5 | 3.8e-17 | 1.9e-15 |
| GO:0044267 | cellular protein metabolic process | 574 | 128 | 71.22 | 1e-13 | 4.4e-12 |

Table S1 – *Continued from previous page*
| GO.ID | Term | Annotated | Significant | Expected | pvalue | FDR |
| --- | --- | --- | --- | --- | --- | --- |
| GO:0019538 | protein metabolic process | 744 | 141 | 92.32 | 2.5e-09 | 9.6e-08 |
| GO:1901564 | organonitrogen compound metabolic proces... | 892 | 161 | 110.68 | 4.5e-09 | 1.5e-07 |
| GO:0034645 | cellular macromolecule biosynthetic proc... | 544 | 106 | 67.5 | 1.3e-07 | 4.0e-06 |
| GO:0009059 | macromolecule biosynthetic process | 546 | 106 | 67.75 | 1.6e-07 | 4.3e-06 |
| GO:0044271 | cellular nitrogen compound biosynthetic ... | 553 | 107 | 68.62 | 1.7e-07 | 4.3e-06 |
| GO:0009058 | biosynthetic process | 700 | 128 | 86.86 | 2.2e-07 | 5.2e-06 |
| GO:0044249 | cellular biosynthetic process | 655 | 121 | 81.28 | 3.1e-07 | 6.8e-06 |
| GO:1901576 | organic substance biosynthetic process | 664 | 122 | 82.39 | 3.7e-07 | 7.6e-06 |
| GO:0009987 | cellular process | 1960 | 288 | 243.21 | 5.8e-07 | 1.1e-5 |
| GO:0044260 | cellular macromolecule metabolic process | 1029 | 170 | 127.68 | 1.4e-06 | 2.5e-5 |
| GO:0044237 | cellular metabolic process | 1425 | 220 | 176.82 | 2.8e-06 | 4.8e-5 |
| GO:0010467 | gene expression | 598 | 107 | 74.2 | 1e-05 | 1.6e-4 |
| GO:0034641 | cellular nitrogen compound metabolic pro... | 872 | 141 | 108.2 | 7.6e-5 | 0.001 |
| GO:0006807 | nitrogen compound metabolic process | 1494 | 216 | 185.38 | 7.0e-4 | 0.010 |
| GO:0008152 | metabolic process | 2076 | 286 | 257.6 | 9.6e-4 | 0.013 |
| GO:0044238 | primary metabolic process | 1584 | 226 | 196.55 | 0.001 | 0.014 |
| GO:0045333 | cellular respiration | 8 | 5 | 0.99 | 0.001 | 0.015 |
| GO:0043170 | macromolecule metabolic process | 1370 | 198 | 170 | 0.002 | 0.020 |
| GO:0046034 | ATP metabolic process | 25 | 9 | 3.1 | 0.002 | 0.025 |
| GO:0071704 | organic substance metabolic process | 1665 | 234 | 206.6 | 0.002 | 0.025 |
| GO:0015980 | energy derivation by oxidation of organi... | 9 | 5 | 1.12 | 0.002 | 0.026 |
| GO:0009144 | purine nucleoside triphosphate metabolic... | 26 | 9 | 3.23 | 0.003 | 0.028 |
| GO:0009199 | ribonucleoside triphosphate metabolic pr... | 26 | 9 | 3.23 | 0.003 | 0.028 |
| GO:0009205 | purine ribonucleoside triphosphate metab... | 26 | 9 | 3.23 | 0.003 | 0.028 |
| GO:0009123 | nucleoside monophosphate metabolic proce... | 27 | 9 | 3.35 | 0.004 | 0.033 |
| GO:0009126 | purine nucleoside monophosphate metaboli... | 27 | 9 | 3.35 | 0.004 | 0.033 |
| GO:0009141 | nucleoside triphosphate metabolic proces... | 27 | 9 | 3.35 | 0.004 | 0.033 |
| GO:0009161 | ribonucleoside monophosphate metabolic p... | 27 | 9 | 3.35 | 0.004 | 0.033 |
| GO:0009167 | purine ribonucleoside monophosphate meta... | 27 | 9 | 3.35 | 0.004 | 0.033 |
| GO:0022900 | electron transport chain | 7 | 4 | 0.87 | 0.006 | 0.050 |
| GO:0006091 | generation of precursor metabolites and ... | 20 | 7 | 2.48 | 0.008 | 0.063 |
| GO:0019693 | ribose phosphate metabolic process | 44 | 11 | 5.46 | 0.016 | 0.122 |
| GO:0007049 | cell cycle | 33 | 9 | 4.09 | 0.016 | 0.122 |
| GO:0009056 | catabolic process | 82 | 17 | 10.18 | 0.021 | 0.149 |
| GO:0006754 | ATP biosynthetic process | 19 | 6 | 2.36 | 0.023 | 0.149 |
| GO:0009142 | nucleoside triphosphate biosynthetic pro... | 19 | 6 | 2.36 | 0.023 | 0.149 |
| GO:0009145 | purine nucleoside triphosphate biosynthe... | 19 | 6 | 2.36 | 0.023 | 0.149 |
| GO:0009201 | ribonucleoside triphosphate biosynthetic... | 19 | 6 | 2.36 | 0.023 | 0.149 |
| GO:0009206 | purine ribonucleoside triphosphate biosy... | 19 | 6 | 2.36 | 0.023 | 0.149 |
| GO:0044257 | cellular protein catabolic process | 35 | 9 | 4.34 | 0.023 | 0.149 |
| GO:0051603 | proteolysis involved in cellular protein... | 35 | 9 | 4.34 | 0.023 | 0.149 |
| GO:0019637 | organophosphate metabolic process | 116 | 22 | 14.39 | 0.025 | 0.159 |
| GO:0007005 | mitochondrion organization | 10 | 4 | 1.24 | 0.027 | 0.163 |
| GO:0044248 | cellular catabolic process | 72 | 15 | 8.93 | 0.028 | 0.165 |
| GO:0006888 | ER to Golgi vesicle-mediated transport | 6 | 3 | 0.74 | 0.028 | 0.165 |
| GO:0009150 | purine ribonucleotide metabolic process | 42 | 10 | 5.21 | 0.029 | 0.165 |
| GO:0009259 | ribonucleotide metabolic process | 42 | 10 | 5.21 | 0.029 | 0.165 |

Table S1 – *Continued from previous page*
| GO.ID | Term | Annotated | Significant | Expected | pvalue | FDR |
| --- | --- | --- | --- | --- | --- | --- |
| GO:0009124 | nucleoside monophosphate biosynthetic pr... | 21 | 6 | 2.61 | 0.037 | 0.193 |
| GO:0009127 | purine nucleoside monophosphate biosynth... | 21 | 6 | 2.61 | 0.037 | 0.193 |
| GO:0009156 | ribonucleoside monophosphate biosyntheti... | 21 | 6 | 2.61 | 0.037 | 0.193 |
| GO:0009168 | purine ribonucleoside monophosphate bios... | 21 | 6 | 2.61 | 0.037 | 0.193 |
| GO:0015985 | energy coupled proton transport, down el... | 11 | 4 | 1.36 | 0.038 | 0.193 |
| GO:0015986 | ATP synthesis coupled proton transport | 11 | 4 | 1.36 | 0.038 | 0.193 |
| GO:0030163 | protein catabolic process | 38 | 9 | 4.72 | 0.039 | 0.194 |
| GO:1901137 | carbohydrate derivative biosynthetic pro... | 76 | 15 | 9.43 | 0.043 | 0.210 |
| GO:1901575 | organic substance catabolic process | 76 | 15 | 9.43 | 0.043 | 0.210 |
| GO:0044265 | cellular macromolecule catabolic process | 45 | 10 | 5.58 | 0.045 | 0.210 |
| GO:0006839 | mitochondrial transport | 7 | 3 | 0.87 | 0.045 | 0.210 |
| GO:1990542 | mitochondrial transmembrane transport | 7 | 3 | 0.87 | 0.045 | 0.210 |
| GO:1901135 | carbohydrate derivative metabolic proces... | 144 | 25 | 17.87 | 0.048 | 0.218 |

**Table S2:**
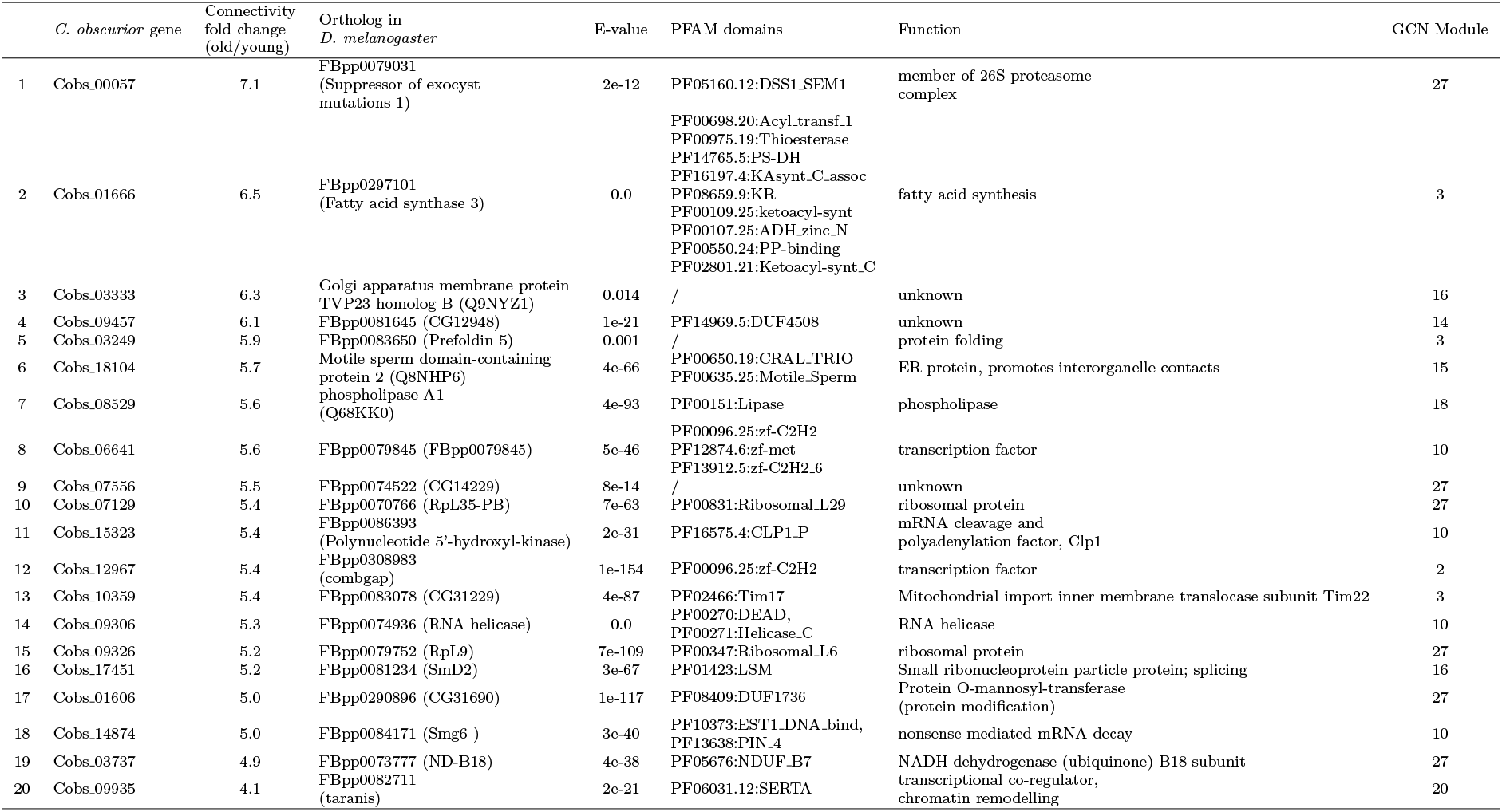
Genes with the greatest increase in connectivity in old compared to young queens.

**Table S3:**
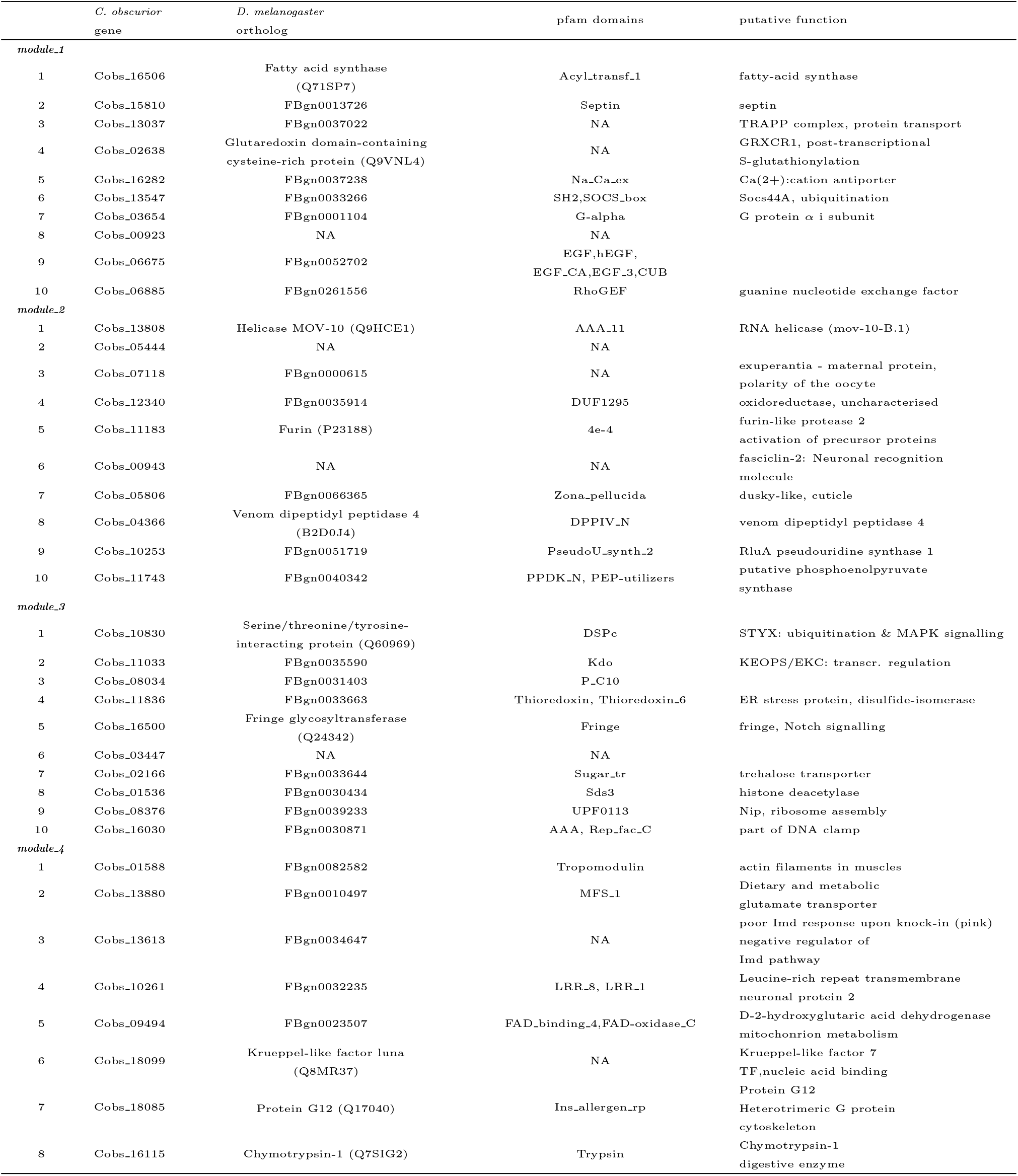
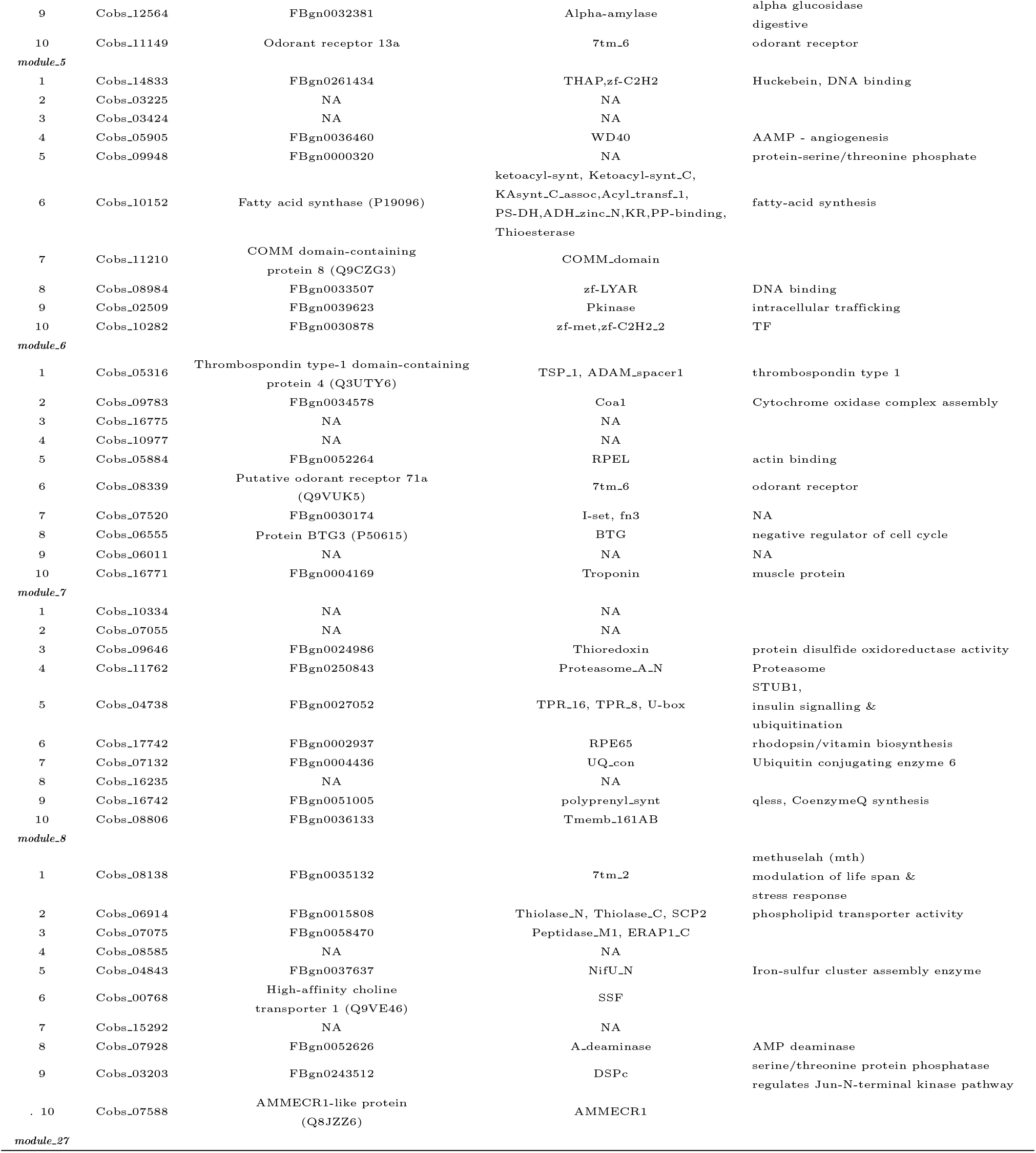
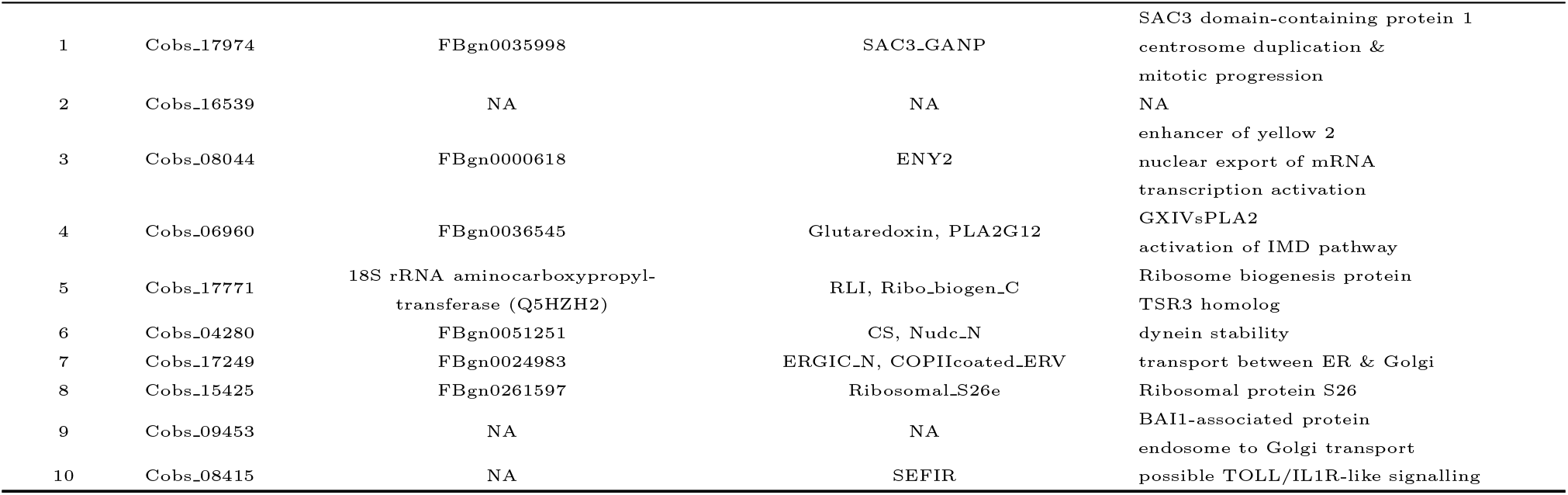
Top hubs of discussed modules within the GCN.

**Table S4:**
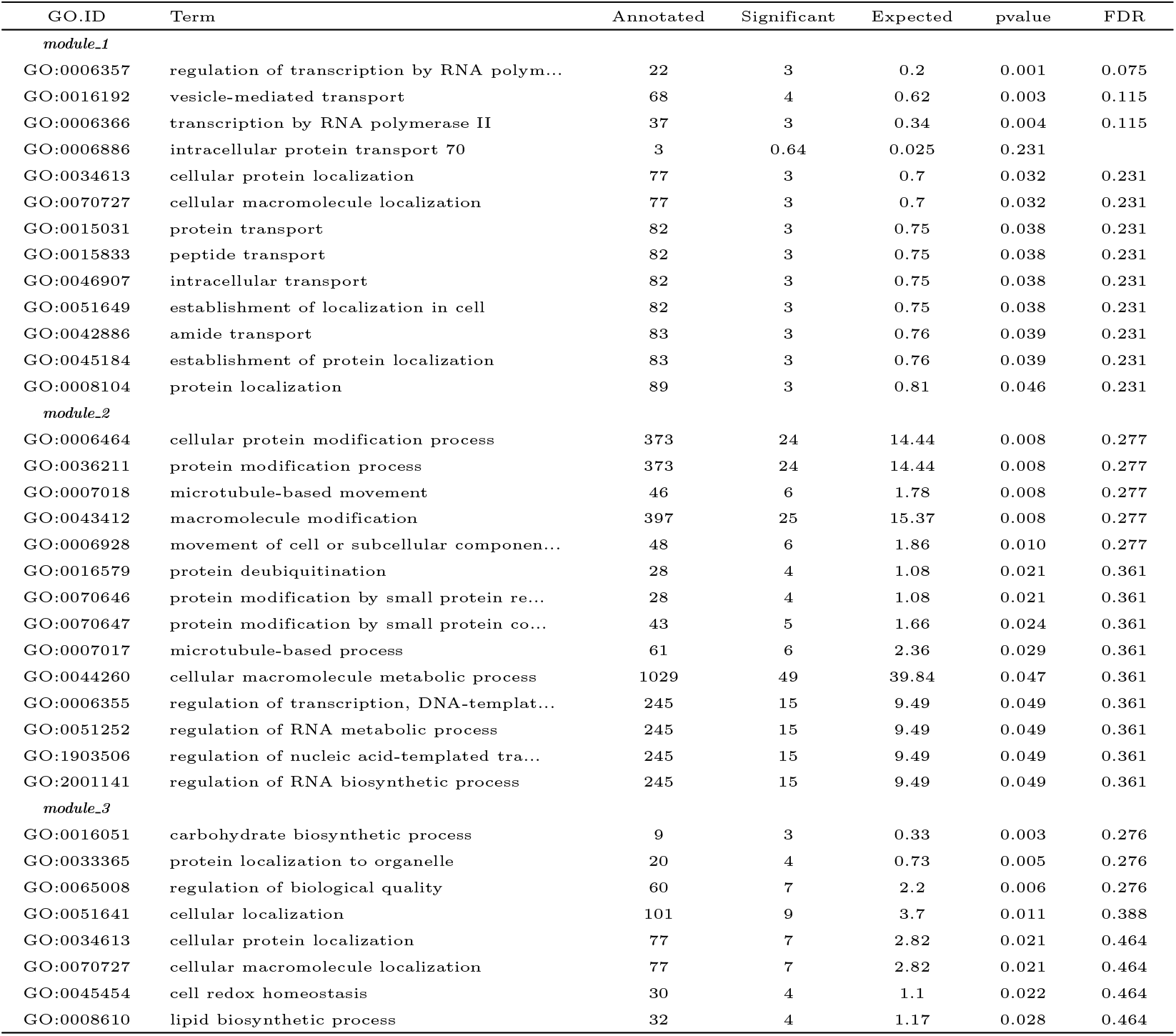
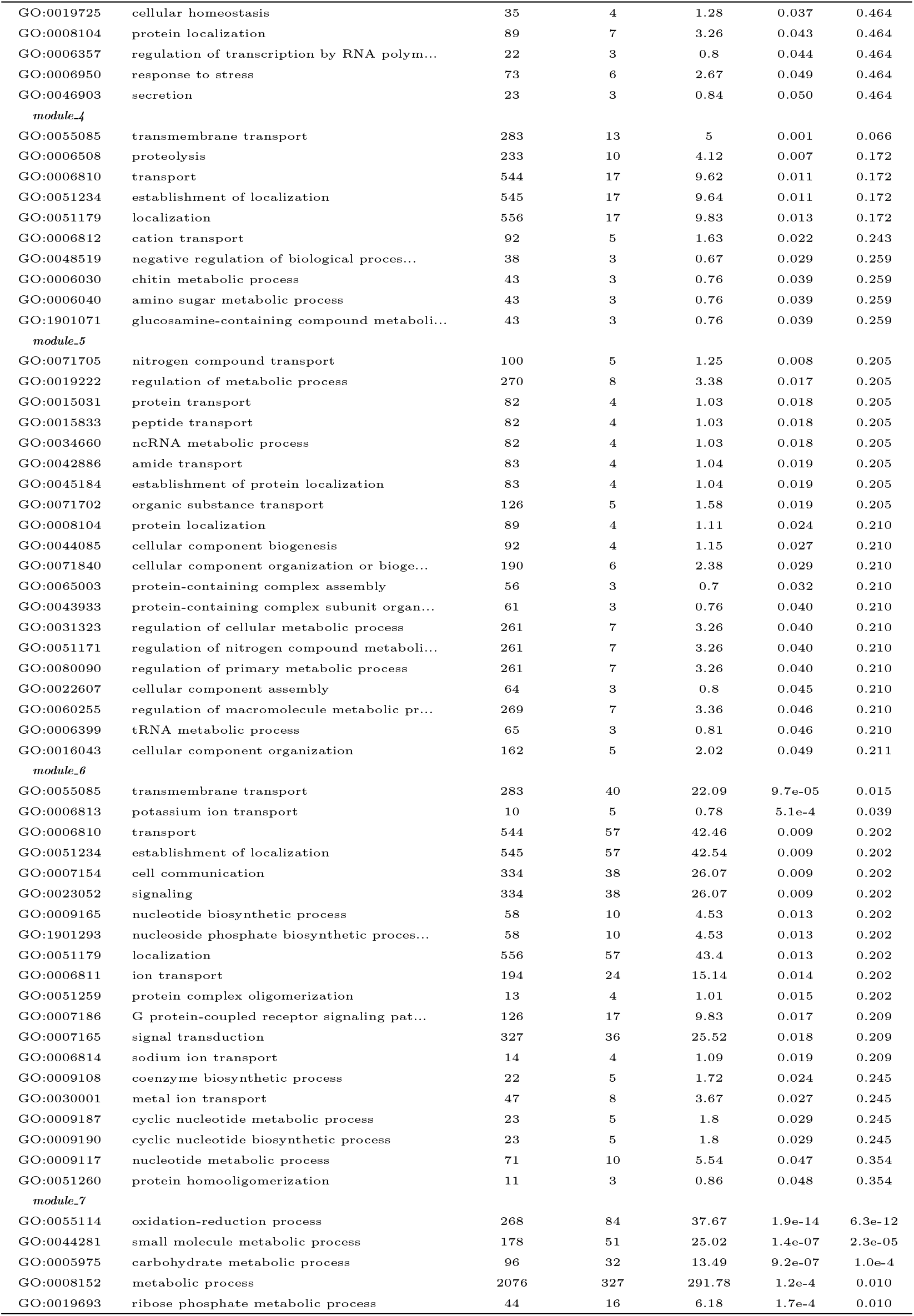
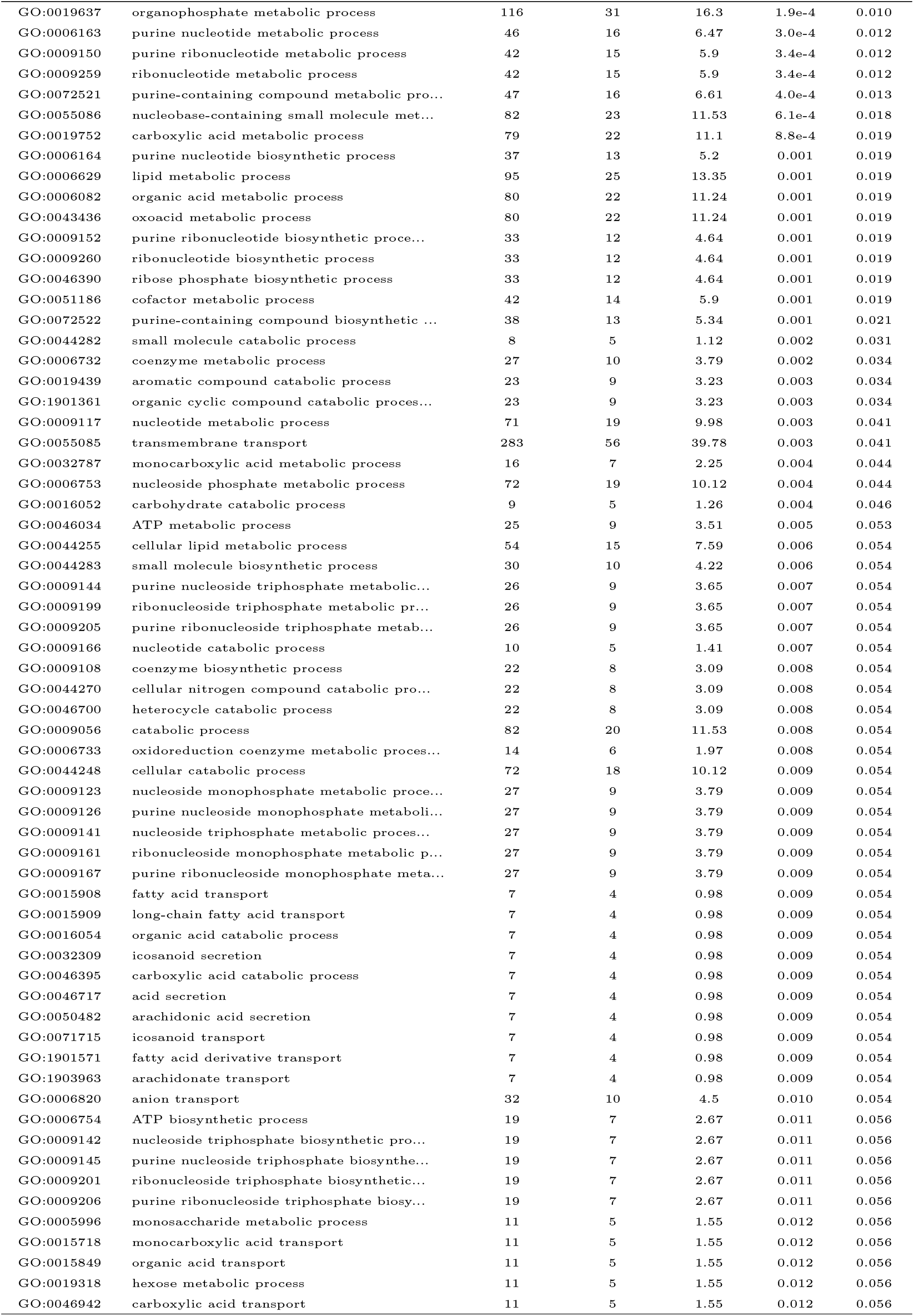
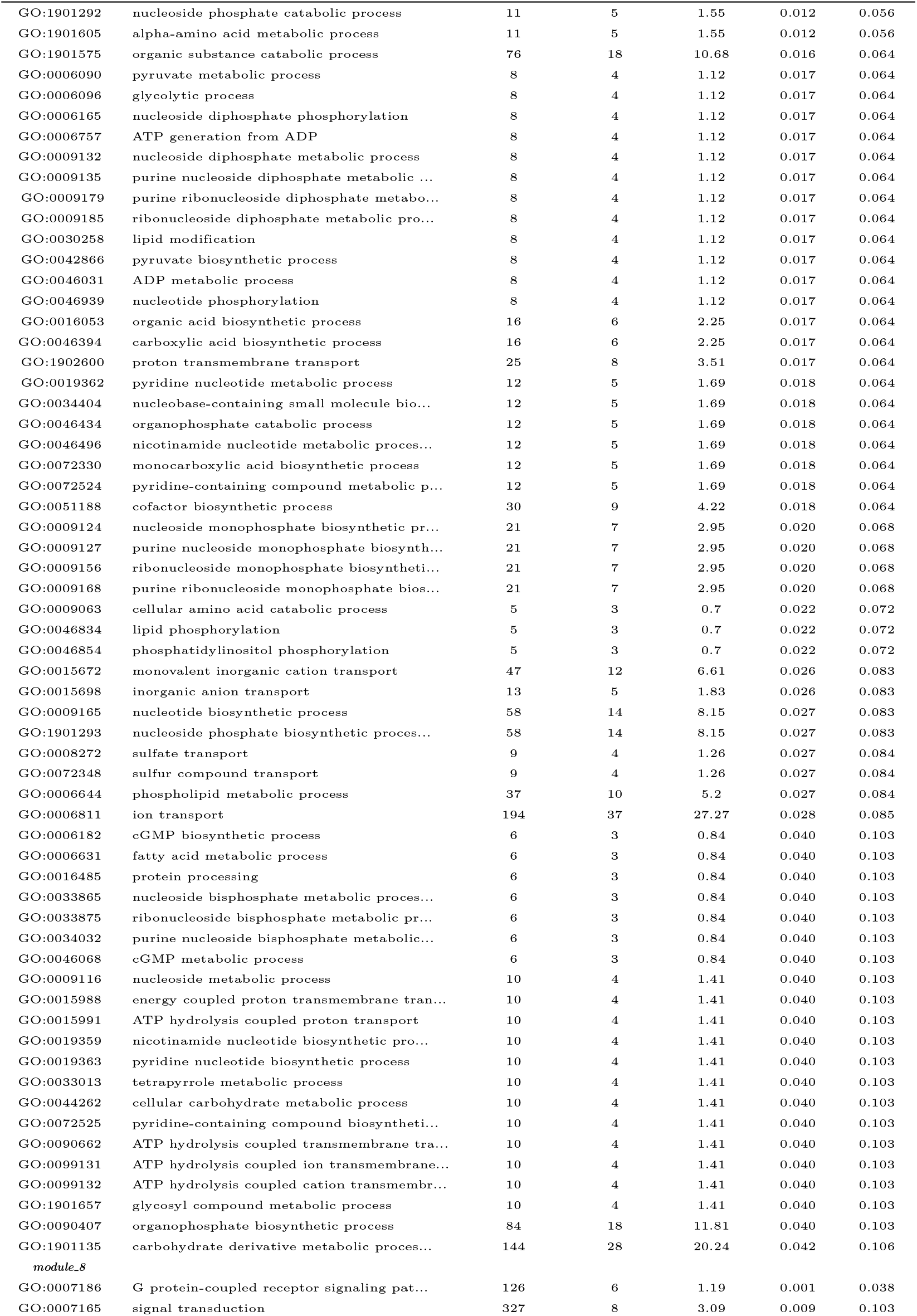
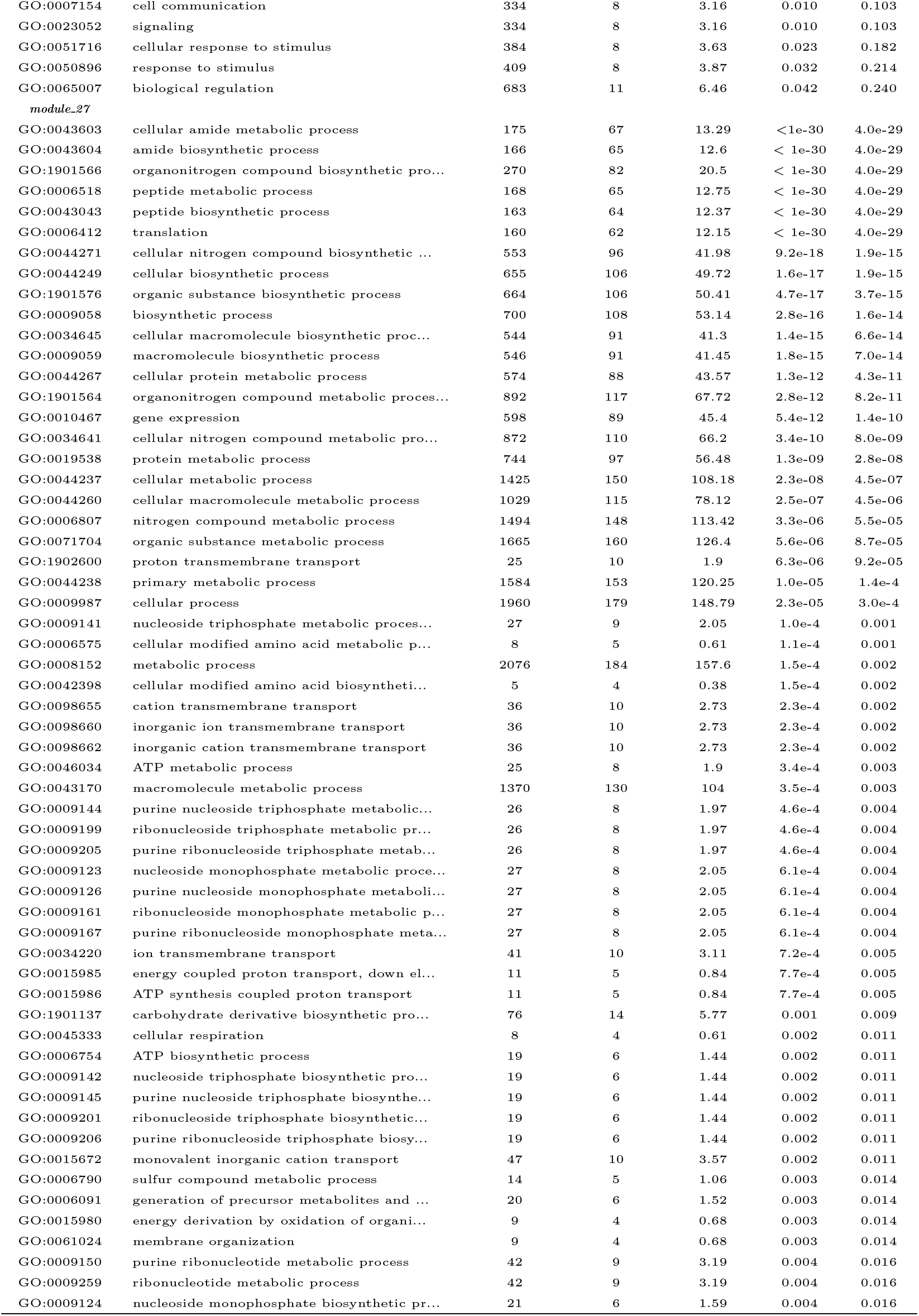
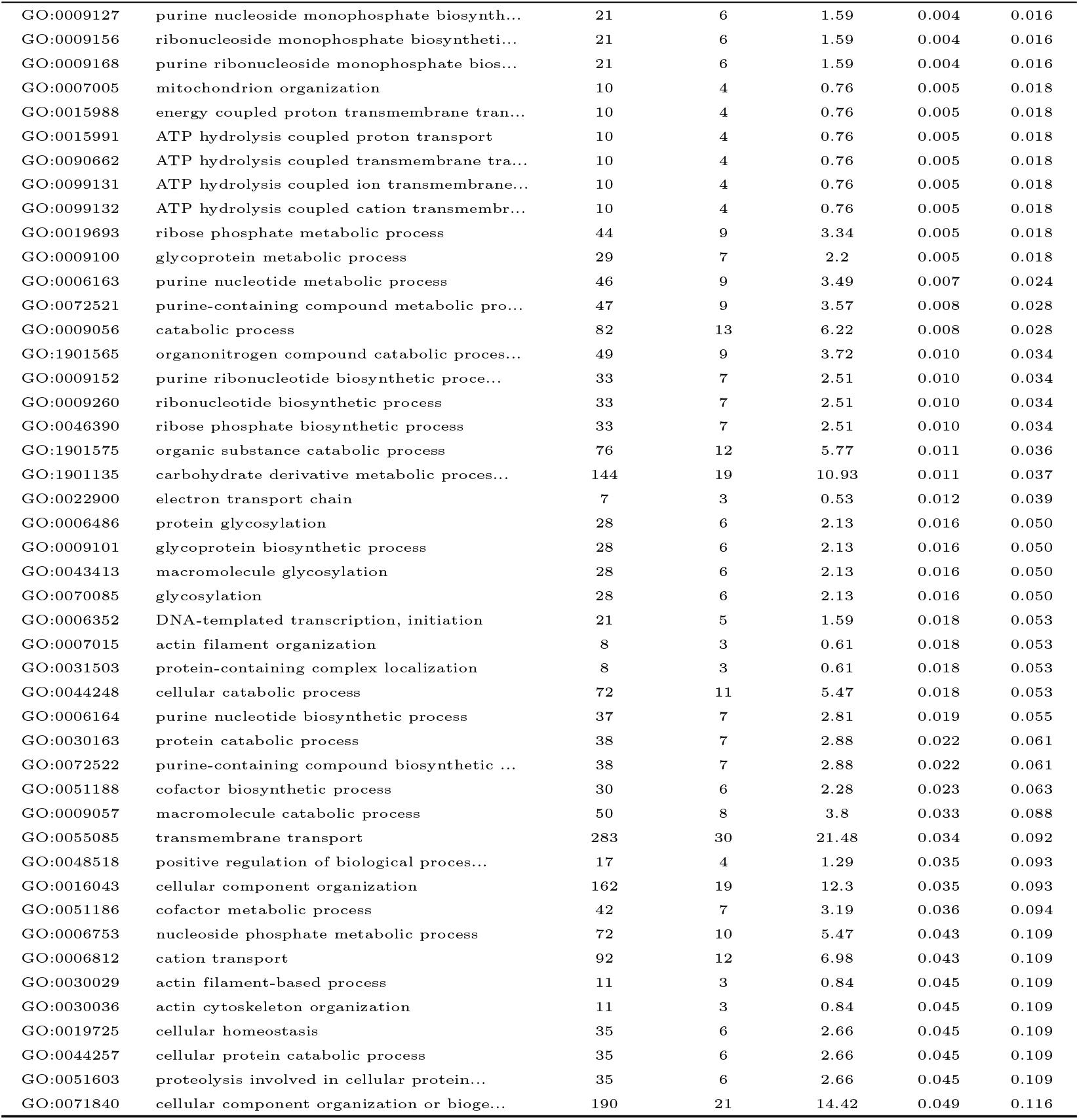
GO terms (Biological Process) significantly enriched within selected modules of the GCN.

| GO.ID | Term | Annotated | Significant | Expected | pvalue | FDR |
| --- | --- | --- | --- | --- | --- | --- |
| *module.1* |  |  |  |  |  |  |
| GO:0006357 | regulation of transcription by RNA polym... | 22 | 3 | 0.2 | 0.001 | 0.075 |
| GO:0016192 | vesicle-mediated transport | 68 | 4 | 0.62 | 0.003 | 0.115 |
| GO:0006366 | transcription by RNA polymerase II | 37 | 3 | 0.34 | 0.004 | 0.115 |
| GO:0006886 | intracellular protein transport 70 | 3 | 0.64 | 0.025 | 0.231 |  |
| GO:0034613 | cellular protein localization | 77 | 3 | 0.7 | 0.032 | 0.231 |
| GO:0070727 | cellular macromolecule localization | 77 | 3 | 0.7 | 0.032 | 0.231 |
| GO:0015031 | protein transport | 82 | 3 | 0.75 | 0.038 | 0.231 |
| GO:0015833 | peptide transport | 82 | 3 | 0.75 | 0.038 | 0.231 |
| GO:0046907 | intracellular transport | 82 | 3 | 0.75 | 0.038 | 0.231 |
| GO:0051649 | establishment of localization in cell | 82 | 3 | 0.75 | 0.038 | 0.231 |
| GO:0042886 | amide transport | 83 | 3 | 0.76 | 0.039 | 0.231 |
| GO:0045184 | establishment of protein localization | 83 | 3 | 0.76 | 0.039 | 0.231 |
| GO:0008104 | protein localization | 89 | 3 | 0.81 | 0.046 | 0.231 |
| *module.2* |  |  |  |  |  |  |
| GO:0006464 | cellular protein modification process | 373 | 24 | 14.44 | 0.008 | 0.277 |
| GO:0036211 | protein modification process | 373 | 24 | 14.44 | 0.008 | 0.277 |
| GO:0007018 | microtubule-based movement | 46 | 6 | 1.78 | 0.008 | 0.277 |
| GO:0043412 | macromolecule modification | 397 | 25 | 15.37 | 0.008 | 0.277 |
| GO:0006928 | movement of cell or subcellular componen... | 48 | 6 | 1.86 | 0.010 | 0.277 |
| GO:0016579 | protein deubiquitination | 28 | 4 | 1.08 | 0.021 | 0.361 |
| GO:0070646 | protein modification by small protein re... | 28 | 4 | 1.08 | 0.021 | 0.361 |
| GO:0070647 | protein modification by small protein co... | 43 | 5 | 1.66 | 0.024 | 0.361 |
| GO:0007017 | microtubule-based process | 61 | 6 | 2.36 | 0.029 | 0.361 |
| GO:0044260 | cellular macromolecule metabolic process | 1029 | 49 | 39.84 | 0.047 | 0.361 |
| GO:0006355 | regulation of transcription, DNA-templat... | 245 | 15 | 9.49 | 0.049 | 0.361 |
| GO:0051252 | regulation of RNA metabolic process | 245 | 15 | 9.49 | 0.049 | 0.361 |
| GO:1903506 | regulation of nucleic acid-templated tra... | 245 | 15 | 9.49 | 0.049 | 0.361 |
| GO:2001141 | regulation of RNA biosynthetic process | 245 | 15 | 9.49 | 0.049 | 0.361 |
| *module.3* |  |  |  |  |  |  |
| GO:0016051 | carbohydrate biosynthetic process | 9 | 3 | 0.33 | 0.003 | 0.276 |
| GO:0033365 | protein localization to organelle | 20 | 4 | 0.73 | 0.005 | 0.276 |
| GO:0065008 | regulation of biological quality | 60 | 7 | 2.2 | 0.006 | 0.276 |
| GO:0051641 | cellular localization | 101 | 9 | 3.7 | 0.011 | 0.388 |
| GO:0034613 | cellular protein localization | 77 | 7 | 2.82 | 0.021 | 0.464 |
| GO:0070727 | cellular macromolecule localization | 77 | 7 | 2.82 | 0.021 | 0.464 |
| GO:0045454 | cell redox homeostasis | 30 | 4 | 1.1 | 0.022 | 0.464 |
| GO:0008610 | lipid biosynthetic process | 32 | 4 | 1.17 | 0.028 | 0.464 |

Table S4 – Continued from previous page
| GO.ID | Term | Annotated | Significant | Expected | pvalue | FDR |
| --- | --- | --- | --- | --- | --- | --- |
| GO:0019725 | cellular homeostasis | 35 | 4 | 1.28 | 0.037 | 0.464 |
| GO:0008104 | protein localization | 89 | 7 | 3.26 | 0.043 | 0.464 |
| GO:0006357 | regulation of transcription by RNA polym... | 22 | 3 | 0.8 | 0.044 | 0.464 |
| GO:0006950 | response to stress | 73 | 6 | 2.67 | 0.049 | 0.464 |
| GO:0046903 | secretion | 23 | 3 | 0.84 | 0.050 | 0.464 |
| *module.4* |  |  |  |  |  |  |
| GO:0055085 | transmembrane transport | 283 | 13 | 5 | 0.001 | 0.066 |
| GO:0006508 | proteolysis | 233 | 10 | 4.12 | 0.007 | 0.172 |
| GO:0006810 | transport | 544 | 17 | 9.62 | 0.011 | 0.172 |
| GO:0051234 | establishment of localization | 545 | 17 | 9.64 | 0.011 | 0.172 |
| GO:0051179 | localization | 556 | 17 | 9.83 | 0.013 | 0.172 |
| GO:0006812 | cation transport | 92 | 5 | 1.63 | 0.022 | 0.243 |
| GO:0048519 | negative regulation of biological proces... | 38 | 3 | 0.67 | 0.029 | 0.259 |
| GO:0006030 | chitin metabolic process | 43 | 3 | 0.76 | 0.039 | 0.259 |
| GO:0006040 | amino sugar metabolic process | 43 | 3 | 0.76 | 0.039 | 0.259 |
| GO:1901071 | glucosamine-containing compound metaboli... | 43 | 3 | 0.76 | 0.039 | 0.259 |
| *module.5* |  |  |  |  |  |  |
| GO:0071705 | nitrogen compound transport | 100 | 5 | 1.25 | 0.008 | 0.205 |
| GO:0019222 | regulation of metabolic process | 270 | 8 | 3.38 | 0.017 | 0.205 |
| GO:0015031 | protein transport | 82 | 4 | 1.03 | 0.018 | 0.205 |
| GO:0015833 | peptide transport | 82 | 4 | 1.03 | 0.018 | 0.205 |
| GO:0034660 | ncRNA metabolic process | 82 | 4 | 1.03 | 0.018 | 0.205 |
| GO:0042886 | amide transport | 83 | 4 | 1.04 | 0.019 | 0.205 |
| GO:0045184 | establishment of protein localization | 83 | 4 | 1.04 | 0.019 | 0.205 |
| GO:0071702 | organic substance transport | 126 | 5 | 1.58 | 0.019 | 0.205 |
| GO:0008104 | protein localization | 89 | 4 | 1.11 | 0.024 | 0.210 |
| GO:0044085 | cellular component biogenesis | 92 | 4 | 1.15 | 0.027 | 0.210 |
| GO:0071840 | cellular component organization or bioge... | 190 | 6 | 2.38 | 0.029 | 0.210 |
| GO:0065003 | protein-containing complex assembly | 56 | 3 | 0.7 | 0.032 | 0.210 |
| GO:0043933 | protein-containing complex subunit organ... | 61 | 3 | 0.76 | 0.040 | 0.210 |
| GO:0031323 | regulation of cellular metabolic process | 261 | 7 | 3.26 | 0.040 | 0.210 |
| GO:0051171 | regulation of nitrogen compound metaboli... | 261 | 7 | 3.26 | 0.040 | 0.210 |
| GO:0080090 | regulation of primary metabolic process | 261 | 7 | 3.26 | 0.040 | 0.210 |
| GO:0022607 | cellular component assembly | 64 | 3 | 0.8 | 0.045 | 0.210 |
| GO:0060255 | regulation of macromolecule metabolic pr... | 269 | 7 | 3.36 | 0.046 | 0.210 |
| GO:0006399 | tRNA metabolic process | 65 | 3 | 0.81 | 0.046 | 0.210 |
| GO:0016043 | cellular component organization | 162 | 5 | 2.02 | 0.049 | 0.211 |
| *module.6* |  |  |  |  |  |  |
| GO:0055085 | transmembrane transport | 283 | 40 | 22.09 | 9.7e-05 | 0.015 |
| GO:0006813 | potassium ion transport | 10 | 5 | 0.78 | 5.1e-4 | 0.039 |
| GO:0006810 | transport | 544 | 57 | 42.46 | 0.009 | 0.202 |
| GO:0051234 | establishment of localization | 545 | 57 | 42.54 | 0.009 | 0.202 |
| GO:0007154 | cell communication | 334 | 38 | 26.07 | 0.009 | 0.202 |
| GO:0023052 | signaling | 334 | 38 | 26.07 | 0.009 | 0.202 |
| GO:0009165 | nucleotide biosynthetic process | 58 | 10 | 4.53 | 0.013 | 0.202 |
| GO:1901293 | nucleoside phosphate biosynthetic proces... | 58 | 10 | 4.53 | 0.013 | 0.202 |
| GO:0051179 | localization | 556 | 57 | 43.4 | 0.013 | 0.202 |
| GO:0006811 | ion transport | 194 | 24 | 15.14 | 0.014 | 0.202 |
| GO:0051259 | protein complex oligomerization | 13 | 4 | 1.01 | 0.015 | 0.202 |
| GO:0007186 | G protein-coupled receptor signaling pat... | 126 | 17 | 9.83 | 0.017 | 0.209 |
| GO:0007165 | signal transduction | 327 | 36 | 25.52 | 0.018 | 0.209 |
| GO:0006814 | sodium ion transport | 14 | 4 | 1.09 | 0.019 | 0.209 |
| GO:0009108 | coenzyme biosynthetic process | 22 | 5 | 1.72 | 0.024 | 0.245 |
| GO:0030001 | metal ion transport | 47 | 8 | 3.67 | 0.027 | 0.245 |
| GO:0009187 | cyclic nucleotide metabolic process | 23 | 5 | 1.8 | 0.029 | 0.245 |
| GO:0009190 | cyclic nucleotide biosynthetic process | 23 | 5 | 1.8 | 0.029 | 0.245 |
| GO:0009117 | nucleotide metabolic process | 71 | 10 | 5.54 | 0.047 | 0.354 |
| GO:0051260 | protein homoooligomerization | 11 | 3 | 0.86 | 0.048 | 0.354 |
| *module.7* |  |  |  |  |  |  |
| GO:0055114 | oxidation-reduction process | 268 | 84 | 37.67 | 1.9e-14 | 6.3e-12 |
| GO:0044281 | small molecule metabolic process | 178 | 51 | 25.02 | 1.4e-07 | 2.3e-05 |
| GO:0005975 | carbohydrate metabolic process | 96 | 32 | 13.49 | 9.2e-07 | 1.0e-4 |
| GO:0008152 | metabolic process | 2076 | 327 | 291.78 | 1.2e-4 | 0.010 |
| GO:0019693 | ribose phosphate metabolic process | 44 | 16 | 6.18 | 1.7e-4 | 0.010 |

Table S4 – Continued from previous page
| GO.ID | Term | Annotated | Significant | Expected | pvalue | FDR |
| --- | --- | --- | --- | --- | --- | --- |
| GO:0019637 | organophosphate metabolic process | 116 | 31 | 16.3 | 1.9e-4 | 0.010 |
| GO:0006163 | purine nucleotide metabolic process | 46 | 16 | 6.47 | 3.0e-4 | 0.012 |
| GO:0009150 | purine ribonucleotide metabolic process | 42 | 15 | 5.9 | 3.4e-4 | 0.012 |
| GO:0009259 | ribonucleotide metabolic process | 42 | 15 | 5.9 | 3.4e-4 | 0.012 |
| GO:0072521 | purine-containing compound metabolic pro... | 47 | 16 | 6.61 | 4.0e-4 | 0.013 |
| GO:0055086 | nucleobase-containing small molecule met... | 82 | 23 | 11.53 | 6.1e-4 | 0.018 |
| GO:0019752 | carboxylic acid metabolic process | 79 | 22 | 11.1 | 8.8e-4 | 0.019 |
| GO:0006164 | purine nucleotide biosynthetic process | 37 | 13 | 5.2 | 0.001 | 0.019 |
| GO:0006629 | lipid metabolic process | 95 | 25 | 13.35 | 0.001 | 0.019 |
| GO:0006082 | organic acid metabolic process | 80 | 22 | 11.24 | 0.001 | 0.019 |
| GO:0043436 | oxoacid metabolic process | 80 | 22 | 11.24 | 0.001 | 0.019 |
| GO:0009152 | purine ribonucleotide biosynthetic proce... | 33 | 12 | 4.64 | 0.001 | 0.019 |
| GO:0009260 | ribonucleotide biosynthetic process | 33 | 12 | 4.64 | 0.001 | 0.019 |
| GO:0046390 | ribose phosphate biosynthetic process | 33 | 12 | 4.64 | 0.001 | 0.019 |
| GO:0051186 | cofactor metabolic process | 42 | 14 | 5.9 | 0.001 | 0.019 |
| GO:0072522 | purine-containing compound biosynthetic ... | 38 | 13 | 5.34 | 0.001 | 0.021 |
| GO:0044282 | small molecule catabolic process | 8 | 5 | 1.12 | 0.002 | 0.031 |
| GO:0006732 | coenzyme metabolic process | 27 | 10 | 3.79 | 0.002 | 0.034 |
| GO:0019439 | aromatic compound catabolic process | 23 | 9 | 3.23 | 0.003 | 0.034 |
| GO:1901361 | organic cyclic compound catabolic proces... | 23 | 9 | 3.23 | 0.003 | 0.034 |
| GO:0009117 | nucleotide metabolic process | 71 | 19 | 9.98 | 0.003 | 0.041 |
| GO:0055085 | transmembrane transport | 283 | 56 | 39.78 | 0.003 | 0.041 |
| GO:0032787 | monocarboxylic acid metabolic process | 16 | 7 | 2.25 | 0.004 | 0.044 |
| GO:0006753 | nucleoside phosphate metabolic process | 72 | 19 | 10.12 | 0.004 | 0.044 |
| GO:0016052 | carbohydrate catabolic process | 9 | 5 | 1.26 | 0.004 | 0.046 |
| GO:0046034 | ATP metabolic process | 25 | 9 | 3.51 | 0.005 | 0.053 |
| GO:0044255 | cellular lipid metabolic process | 54 | 15 | 7.59 | 0.006 | 0.054 |
| GO:0044283 | small molecule biosynthetic process | 30 | 10 | 4.22 | 0.006 | 0.054 |
| GO:0009144 | purine nucleoside triphosphate metabolic... | 26 | 9 | 3.65 | 0.007 | 0.054 |
| GO:0009199 | ribonucleoside triphosphate metabolic pr... | 26 | 9 | 3.65 | 0.007 | 0.054 |
| GO:0009205 | purine ribonucleoside triphosphate metab... | 26 | 9 | 3.65 | 0.007 | 0.054 |
| GO:0009166 | nucleotide catabolic process | 10 | 5 | 1.41 | 0.007 | 0.054 |
| GO:0009108 | coenzyme biosynthetic process | 22 | 8 | 3.09 | 0.008 | 0.054 |
| GO:0044270 | cellular nitrogen compound catabolic pro... | 22 | 8 | 3.09 | 0.008 | 0.054 |
| GO:0046700 | heterocycle catabolic process | 22 | 8 | 3.09 | 0.008 | 0.054 |
| GO:0009056 | catabolic process | 82 | 20 | 11.53 | 0.008 | 0.054 |
| GO:0006733 | oxidoreduction coenzyme metabolic proces... | 14 | 6 | 1.97 | 0.008 | 0.054 |
| GO:0044248 | cellular catabolic process | 72 | 18 | 10.12 | 0.009 | 0.054 |
| GO:0009123 | nucleoside monophosphate metabolic proce... | 27 | 9 | 3.79 | 0.009 | 0.054 |
| GO:0009126 | purine nucleoside monophosphate metaboli... | 27 | 9 | 3.79 | 0.009 | 0.054 |
| GO:0009141 | nucleoside triphosphate metabolic proces... | 27 | 9 | 3.79 | 0.009 | 0.054 |
| GO:0009161 | ribonucleoside monophosphate metabolic p... | 27 | 9 | 3.79 | 0.009 | 0.054 |
| GO:0009167 | purine ribonucleoside monophosphate meta... | 27 | 9 | 3.79 | 0.009 | 0.054 |
| GO:0015908 | fatty acid transport | 7 | 4 | 0.98 | 0.009 | 0.054 |
| GO:0015909 | long-chain fatty acid transport | 7 | 4 | 0.98 | 0.009 | 0.054 |
| GO:0016054 | organic acid catabolic process | 7 | 4 | 0.98 | 0.009 | 0.054 |
| GO:0032309 | icosanoid secretion | 7 | 4 | 0.98 | 0.009 | 0.054 |
| GO:0046395 | carboxylic acid catabolic process | 7 | 4 | 0.98 | 0.009 | 0.054 |
| GO:0046717 | acid secretion | 7 | 4 | 0.98 | 0.009 | 0.054 |
| GO:0050482 | arachidonic acid secretion | 7 | 4 | 0.98 | 0.009 | 0.054 |
| GO:0071715 | icosanoid transport | 7 | 4 | 0.98 | 0.009 | 0.054 |
| GO:1901571 | fatty acid derivative transport | 7 | 4 | 0.98 | 0.009 | 0.054 |
| GO:1903963 | arachidonate transport | 7 | 4 | 0.98 | 0.009 | 0.054 |
| GO:0006820 | anion transport | 32 | 10 | 4.5 | 0.010 | 0.054 |
| GO:0006754 | ATP biosynthetic process | 19 | 7 | 2.67 | 0.011 | 0.056 |
| GO:0009142 | nucleoside triphosphate biosynthetic pro... | 19 | 7 | 2.67 | 0.011 | 0.056 |
| GO:0009145 | purine nucleoside triphosphate biosynthe... | 19 | 7 | 2.67 | 0.011 | 0.056 |
| GO:0009201 | ribonucleoside triphosphate biosynthetic... | 19 | 7 | 2.67 | 0.011 | 0.056 |
| GO:0009206 | purine ribonucleoside triphosphate biosy... | 19 | 7 | 2.67 | 0.011 | 0.056 |
| GO:0005996 | monosaccharide metabolic process | 11 | 5 | 1.55 | 0.012 | 0.056 |
| GO:0015718 | monocarboxylic acid transport | 11 | 5 | 1.55 | 0.012 | 0.056 |
| GO:0015849 | organic acid transport | 11 | 5 | 1.55 | 0.012 | 0.056 |
| GO:0019318 | hexose metabolic process | 11 | 5 | 1.55 | 0.012 | 0.056 |
| GO:0046942 | carboxylic acid transport | 11 | 5 | 1.55 | 0.012 | 0.056 |

Table S4 – Continued from previous page
| GO.ID | Term | Annotated | Significant | Expected | pvalue | FDR |
| --- | --- | --- | --- | --- | --- | --- |
| GO:1901292 | nucleoside phosphate catabolic process | 11 | 5 | 1.55 | 0.012 | 0.056 |
| GO:1901605 | alpha-amino acid metabolic process | 11 | 5 | 1.55 | 0.012 | 0.056 |
| GO:1901575 | organic substance catabolic process | 76 | 18 | 10.68 | 0.016 | 0.064 |
| GO:0006090 | pyruvate metabolic process | 8 | 4 | 1.12 | 0.017 | 0.064 |
| GO:0006096 | glycolytic process | 8 | 4 | 1.12 | 0.017 | 0.064 |
| GO:0006165 | nucleoside diphosphate phosphorylation | 8 | 4 | 1.12 | 0.017 | 0.064 |
| GO:0006757 | ATP generation from ADP | 8 | 4 | 1.12 | 0.017 | 0.064 |
| GO:0009132 | nucleoside diphosphate metabolic process | 8 | 4 | 1.12 | 0.017 | 0.064 |
| GO:0009135 | purine nucleoside diphosphate metabolic ... | 8 | 4 | 1.12 | 0.017 | 0.064 |
| GO:0009179 | purine ribonucleoside diphosphate metabo... | 8 | 4 | 1.12 | 0.017 | 0.064 |
| GO:0009185 | ribonucleoside diphosphate metabolic pro... | 8 | 4 | 1.12 | 0.017 | 0.064 |
| GO:0030258 | lipid modification | 8 | 4 | 1.12 | 0.017 | 0.064 |
| GO:0042866 | pyruvate biosynthetic process | 8 | 4 | 1.12 | 0.017 | 0.064 |
| GO:0046031 | ADP metabolic process | 8 | 4 | 1.12 | 0.017 | 0.064 |
| GO:0046939 | nucleotide phosphorylation | 8 | 4 | 1.12 | 0.017 | 0.064 |
| GO:0016053 | organic acid biosynthetic process | 16 | 6 | 2.25 | 0.017 | 0.064 |
| GO:0046394 | carboxylic acid biosynthetic process | 16 | 6 | 2.25 | 0.017 | 0.064 |
| GO:1902600 | proton transmembrane transport | 25 | 8 | 3.51 | 0.017 | 0.064 |
| GO:0019362 | pyridine nucleotide metabolic process | 12 | 5 | 1.69 | 0.018 | 0.064 |
| GO:0034404 | nucleobase-containing small molecule bio... | 12 | 5 | 1.69 | 0.018 | 0.064 |
| GO:0046434 | organophosphate catabolic process | 12 | 5 | 1.69 | 0.018 | 0.064 |
| GO:0046496 | nicotinamide nucleotide metabolic proces... | 12 | 5 | 1.69 | 0.018 | 0.064 |
| GO:0072330 | monocarboxylic acid biosynthetic process | 12 | 5 | 1.69 | 0.018 | 0.064 |
| GO:0072524 | pyridine-containing compound metabolic p... | 12 | 5 | 1.69 | 0.018 | 0.064 |
| GO:0051188 | cofactor biosynthetic process | 30 | 9 | 4.22 | 0.018 | 0.064 |
| GO:0009124 | nucleoside monophosphate biosynthetic pr... | 21 | 7 | 2.95 | 0.020 | 0.068 |
| GO:0009127 | purine nucleoside monophosphate biosynth... | 21 | 7 | 2.95 | 0.020 | 0.068 |
| GO:0009156 | ribonucleoside monophosphate biosyntheti... | 21 | 7 | 2.95 | 0.020 | 0.068 |
| GO:0009168 | purine ribonucleoside monophosphate bios... | 21 | 7 | 2.95 | 0.020 | 0.068 |
| GO:0009063 | cellular amino acid catabolic process | 5 | 3 | 0.7 | 0.022 | 0.072 |
| GO:0046834 | lipid phosphorylation | 5 | 3 | 0.7 | 0.022 | 0.072 |
| GO:0046854 | phosphatidylinositol phosphorylation | 5 | 3 | 0.7 | 0.022 | 0.072 |
| GO:0015672 | monovalent inorganic cation transport | 47 | 12 | 6.61 | 0.026 | 0.083 |
| GO:0015698 | inorganic anion transport | 13 | 5 | 1.83 | 0.026 | 0.083 |
| GO:0009165 | nucleotide biosynthetic process | 58 | 14 | 8.15 | 0.027 | 0.083 |
| GO:1901293 | nucleoside phosphate biosynthetic proces... | 58 | 14 | 8.15 | 0.027 | 0.083 |
| GO:0008272 | sulfate transport | 9 | 4 | 1.26 | 0.027 | 0.084 |
| GO:0072348 | sulfur compound transport | 9 | 4 | 1.26 | 0.027 | 0.084 |
| GO:0006644 | phospholipid metabolic process | 37 | 10 | 5.2 | 0.027 | 0.084 |
| GO:0006811 | ion transport | 194 | 37 | 27.27 | 0.028 | 0.085 |
| GO:0006182 | cGMP biosynthetic process | 6 | 3 | 0.84 | 0.040 | 0.103 |
| GO:0006631 | fatty acid metabolic process | 6 | 3 | 0.84 | 0.040 | 0.103 |
| GO:0016485 | protein processing | 6 | 3 | 0.84 | 0.040 | 0.103 |
| GO:0033865 | nucleoside bisphosphate metabolic proces... | 6 | 3 | 0.84 | 0.040 | 0.103 |
| GO:0033875 | ribonucleoside bisphosphate metabolic pr... | 6 | 3 | 0.84 | 0.040 | 0.103 |
| GO:0034032 | purine nucleoside bisphosphate metabolic... | 6 | 3 | 0.84 | 0.040 | 0.103 |
| GO:0046068 | cGMP metabolic process | 6 | 3 | 0.84 | 0.040 | 0.103 |
| GO:0009116 | nucleoside metabolic process | 10 | 4 | 1.41 | 0.040 | 0.103 |
| GO:0015988 | energy coupled proton transmembrane tran... | 10 | 4 | 1.41 | 0.040 | 0.103 |
| GO:0015991 | ATP hydrolysis coupled proton transport | 10 | 4 | 1.41 | 0.040 | 0.103 |
| GO:0019359 | nicotinamide nucleotide biosynthetic pro... | 10 | 4 | 1.41 | 0.040 | 0.103 |
| GO:0019363 | pyridine nucleotide biosynthetic process | 10 | 4 | 1.41 | 0.040 | 0.103 |
| GO:0033013 | tetrapyrrole metabolic process | 10 | 4 | 1.41 | 0.040 | 0.103 |
| GO:0044262 | cellular carbohydrate metabolic process | 10 | 4 | 1.41 | 0.040 | 0.103 |
| GO:0072525 | pyridine-containing compound biosyntheti... | 10 | 4 | 1.41 | 0.040 | 0.103 |
| GO:0090662 | ATP hydrolysis coupled transmembrane tra... | 10 | 4 | 1.41 | 0.040 | 0.103 |
| GO:0099131 | ATP hydrolysis coupled ion transmembrane... | 10 | 4 | 1.41 | 0.040 | 0.103 |
| GO:0099132 | ATP hydrolysis coupled cation transmembr... | 10 | 4 | 1.41 | 0.040 | 0.103 |
| GO:1901657 | glycosyl compound metabolic process | 10 | 4 | 1.41 | 0.040 | 0.103 |
| GO:0090407 | organophosphate biosynthetic process | 84 | 18 | 11.81 | 0.040 | 0.103 |
| GO:1901135 | carbohydrate derivative metabolic proces... | 144 | 28 | 20.24 | 0.042 | 0.106 |
| *module_8* |  |  |  |  |  |  |
| GO:0007186 | G protein-coupled receptor signaling pat... | 126 | 6 | 1.19 | 0.001 | 0.038 |
| GO:0007165 | signal transduction | 327 | 8 | 3.09 | 0.009 | 0.103 |

Table S4 – Continued from previous page
| GO.ID | Term | Annotated | Significant | Expected | pvalue | FDR |
| --- | --- | --- | --- | --- | --- | --- |
| GO:0007154 | cell communication | 334 | 8 | 3.16 | 0.010 | 0.103 |
| GO:0023052 | signaling | 334 | 8 | 3.16 | 0.010 | 0.103 |
| GO:0051716 | cellular response to stimulus | 384 | 8 | 3.63 | 0.023 | 0.182 |
| GO:0050896 | response to stimulus | 409 | 8 | 3.87 | 0.032 | 0.214 |
| GO:0065007 | biological regulation | 683 | 11 | 6.46 | 0.042 | 0.240 |
| *module_27* |  |  |  |  |  |  |
| GO:0043603 | cellular amide metabolic process | 175 | 67 | 13.29 | <1e-30 | 4.0e-29 |
| GO:0043604 | amide biosynthetic process | 166 | 65 | 12.6 | < 1e-30 | 4.0e-29 |
| GO:1901566 | organonitrogen compound biosynthetic pro... | 270 | 82 | 20.5 | < 1e-30 | 4.0e-29 |
| GO:0006518 | peptide metabolic process | 168 | 65 | 12.75 | < 1e-30 | 4.0e-29 |
| GO:0043043 | peptide biosynthetic process | 163 | 64 | 12.37 | < 1e-30 | 4.0e-29 |
| GO:0006412 | translation | 160 | 62 | 12.15 | < 1e-30 | 4.0e-29 |
| GO:0044271 | cellular nitrogen compound biosynthetic ... | 553 | 96 | 41.98 | 9.2e-18 | 1.9e-15 |
| GO:0044249 | cellular biosynthetic process | 655 | 106 | 49.72 | 1.6e-17 | 1.9e-15 |
| GO:1901576 | organic substance biosynthetic process | 664 | 106 | 50.41 | 4.7e-17 | 3.7e-15 |
| GO:0009058 | biosynthetic process | 700 | 108 | 53.14 | 2.8e-16 | 1.6e-14 |
| GO:0034645 | cellular macromolecule biosynthetic proc... | 544 | 91 | 41.3 | 1.4e-15 | 6.6e-14 |
| GO:0009059 | macromolecule biosynthetic process | 546 | 91 | 41.45 | 1.8e-15 | 7.0e-14 |
| GO:0044267 | cellular protein metabolic process | 574 | 88 | 43.57 | 1.3e-12 | 4.3e-11 |
| GO:1901564 | organonitrogen compound metabolic proces... | 892 | 117 | 67.72 | 2.8e-12 | 8.2e-11 |
| GO:0010467 | gene expression | 598 | 89 | 45.4 | 5.4e-12 | 1.4e-10 |
| GO:0034641 | cellular nitrogen compound metabolic pro... | 872 | 110 | 66.2 | 3.4e-10 | 8.0e-09 |
| GO:0019538 | protein metabolic process | 744 | 97 | 56.48 | 1.3e-09 | 2.8e-08 |
| GO:0044237 | cellular metabolic process | 1425 | 150 | 108.18 | 2.3e-08 | 4.5e-07 |
| GO:0044260 | cellular macromolecule metabolic process | 1029 | 115 | 78.12 | 2.5e-07 | 4.5e-06 |
| GO:0006807 | nitrogen compound metabolic process | 1494 | 148 | 113.42 | 3.3e-06 | 5.5e-05 |
| GO:0071704 | organic substance metabolic process | 1665 | 160 | 126.4 | 5.6e-06 | 8.7e-05 |
| GO:1902600 | proton transmembrane transport | 25 | 10 | 1.9 | 6.3e-06 | 9.2e-05 |
| GO:0044238 | primary metabolic process | 1584 | 153 | 120.25 | 1.0e-05 | 1.4e-4 |
| GO:0009987 | cellular process | 1960 | 179 | 148.79 | 2.3e-05 | 3.0e-4 |
| GO:0009141 | nucleoside triphosphate metabolic proces... | 27 | 9 | 2.05 | 1.0e-4 | 0.001 |
| GO:0006575 | cellular modified amino acid metabolic p... | 8 | 5 | 0.61 | 1.1e-4 | 0.001 |
| GO:0008152 | metabolic process | 2076 | 184 | 157.6 | 1.5e-4 | 0.002 |
| GO:0042398 | cellular modified amino acid biosyntheti... | 5 | 4 | 0.38 | 1.5e-4 | 0.002 |
| GO:0098655 | cation transmembrane transport | 36 | 10 | 2.73 | 2.3e-4 | 0.002 |
| GO:0098660 | inorganic ion transmembrane transport | 36 | 10 | 2.73 | 2.3e-4 | 0.002 |
| GO:0098662 | inorganic cation transmembrane transport | 36 | 10 | 2.73 | 2.3e-4 | 0.002 |
| GO:0046034 | ATP metabolic process | 25 | 8 | 1.9 | 3.4e-4 | 0.003 |
| GO:0043170 | macromolecule metabolic process | 1370 | 130 | 104 | 3.5e-4 | 0.003 |
| GO:0009144 | purine nucleoside triphosphate metabolic... | 26 | 8 | 1.97 | 4.6e-4 | 0.004 |
| GO:0009199 | ribonucleoside triphosphate metabolic pr... | 26 | 8 | 1.97 | 4.6e-4 | 0.004 |
| GO:0009205 | purine ribonucleoside triphosphate metab... | 26 | 8 | 1.97 | 4.6e-4 | 0.004 |
| GO:0009123 | nucleoside monophosphate metabolic proce... | 27 | 8 | 2.05 | 6.1e-4 | 0.004 |
| GO:0009126 | purine nucleoside monophosphate metaboli... | 27 | 8 | 2.05 | 6.1e-4 | 0.004 |
| GO:0009161 | ribonucleoside monophosphate metabolic p... | 27 | 8 | 2.05 | 6.1e-4 | 0.004 |
| GO:0009167 | purine ribonucleoside monophosphate meta... | 27 | 8 | 2.05 | 6.1e-4 | 0.004 |
| GO:0034220 | ion transmembrane transport | 41 | 10 | 3.11 | 7.2e-4 | 0.005 |
| GO:0015985 | energy coupled proton transport, down el... | 11 | 5 | 0.84 | 7.7e-4 | 0.005 |
| GO:0015986 | ATP synthesis coupled proton transport | 11 | 5 | 0.84 | 7.7e-4 | 0.005 |
| GO:1901137 | carbohydrate derivative biosynthetic pro... | 76 | 14 | 5.77 | 0.001 | 0.009 |
| GO:0045333 | cellular respiration | 8 | 4 | 0.61 | 0.002 | 0.011 |
| GO:0006754 | ATP biosynthetic process | 19 | 6 | 1.44 | 0.002 | 0.011 |
| GO:0009142 | nucleoside triphosphate biosynthetic pro... | 19 | 6 | 1.44 | 0.002 | 0.011 |
| GO:0009145 | purine nucleoside triphosphate biosynthe... | 19 | 6 | 1.44 | 0.002 | 0.011 |
| GO:0009201 | ribonucleoside triphosphate biosynthetic... | 19 | 6 | 1.44 | 0.002 | 0.011 |
| GO:0009206 | purine ribonucleoside triphosphate biosy... | 19 | 6 | 1.44 | 0.002 | 0.011 |
| GO:0015672 | monovalent inorganic cation transport | 47 | 10 | 3.57 | 0.002 | 0.011 |
| GO:0006790 | sulfur compound metabolic process | 14 | 5 | 1.06 | 0.003 | 0.014 |
| GO:0006091 | generation of precursor metabolites and ... | 20 | 6 | 1.52 | 0.003 | 0.014 |
| GO:0015980 | energy derivation by oxidation of organi... | 9 | 4 | 0.68 | 0.003 | 0.014 |
| GO:0061024 | membrane organization | 9 | 4 | 0.68 | 0.003 | 0.014 |
| GO:0009150 | purine ribonucleotide metabolic process | 42 | 9 | 3.19 | 0.004 | 0.016 |
| GO:0009259 | ribonucleotide metabolic process | 42 | 9 | 3.19 | 0.004 | 0.016 |
| GO:0009124 | nucleoside monophosphate biosynthetic pr... | 21 | 6 | 1.59 | 0.004 | 0.016 |

Table S4 – Continued from previous page
| GO.ID | Term | Annotated | Significant | Expected | pvalue | FDR |
| --- | --- | --- | --- | --- | --- | --- |
| GO:0009127 | purine nucleoside monophosphate biosynth... | 21 | 6 | 1.59 | 0.004 | 0.016 |
| GO:0009156 | ribonucleoside monophosphate biosyntheti... | 21 | 6 | 1.59 | 0.004 | 0.016 |
| GO:0009168 | purine ribonucleoside monophosphate bios... | 21 | 6 | 1.59 | 0.004 | 0.016 |
| GO:0007005 | mitochondrion organization | 10 | 4 | 0.76 | 0.005 | 0.018 |
| GO:0015988 | energy coupled proton transmembrane tran... | 10 | 4 | 0.76 | 0.005 | 0.018 |
| GO:0015991 | ATP hydrolysis coupled proton transport | 10 | 4 | 0.76 | 0.005 | 0.018 |
| GO:0090662 | ATP hydrolysis coupled transmembrane tra... | 10 | 4 | 0.76 | 0.005 | 0.018 |
| GO:0099131 | ATP hydrolysis coupled ion transmembrane... | 10 | 4 | 0.76 | 0.005 | 0.018 |
| GO:0099132 | ATP hydrolysis coupled cation transmembr... | 10 | 4 | 0.76 | 0.005 | 0.018 |
| GO:0019693 | ribose phosphate metabolic process | 44 | 9 | 3.34 | 0.005 | 0.018 |
| GO:0009100 | glycoprotein metabolic process | 29 | 7 | 2.2 | 0.005 | 0.018 |
| GO:0006163 | purine nucleotide metabolic process | 46 | 9 | 3.49 | 0.007 | 0.024 |
| GO:0072521 | purine-containing compound metabolic pro... | 47 | 9 | 3.57 | 0.008 | 0.028 |
| GO:0009056 | catabolic process | 82 | 13 | 6.22 | 0.008 | 0.028 |
| GO:1901565 | organonitrogen compound catabolic proces... | 49 | 9 | 3.72 | 0.010 | 0.034 |
| GO:0009152 | purine ribonucleotide biosynthetic proces... | 33 | 7 | 2.51 | 0.010 | 0.034 |
| GO:0009260 | ribonucleotide biosynthetic process | 33 | 7 | 2.51 | 0.010 | 0.034 |
| GO:0046390 | ribose phosphate biosynthetic process | 33 | 7 | 2.51 | 0.010 | 0.034 |
| GO:1901575 | organic substance catabolic process | 76 | 12 | 5.77 | 0.011 | 0.036 |
| GO:1901135 | carbohydrate derivative metabolic proces... | 144 | 19 | 10.93 | 0.011 | 0.037 |
| GO:0022900 | electron transport chain | 7 | 3 | 0.53 | 0.012 | 0.039 |
| GO:0006486 | protein glycosylation | 28 | 6 | 2.13 | 0.016 | 0.050 |
| GO:0009101 | glycoprotein biosynthetic process | 28 | 6 | 2.13 | 0.016 | 0.050 |
| GO:0043413 | macromolecule glycosylation | 28 | 6 | 2.13 | 0.016 | 0.050 |
| GO:0070085 | glycosylation | 28 | 6 | 2.13 | 0.016 | 0.050 |
| GO:0006352 | DNA-templated transcription, initiation | 21 | 5 | 1.59 | 0.018 | 0.053 |
| GO:0007015 | actin filament organization | 8 | 3 | 0.61 | 0.018 | 0.053 |
| GO:0031503 | protein-containing complex localization | 8 | 3 | 0.61 | 0.018 | 0.053 |
| GO:0044248 | cellular catabolic process | 72 | 11 | 5.47 | 0.018 | 0.053 |
| GO:0006164 | purine nucleotide biosynthetic process | 37 | 7 | 2.81 | 0.019 | 0.055 |
| GO:0030163 | protein catabolic process | 38 | 7 | 2.88 | 0.022 | 0.061 |
| GO:0072522 | purine-containing compound biosynthetic ... | 38 | 7 | 2.88 | 0.022 | 0.061 |
| GO:0051188 | cofactor biosynthetic process | 30 | 6 | 2.28 | 0.023 | 0.063 |
| GO:0009057 | macromolecule catabolic process | 50 | 8 | 3.8 | 0.033 | 0.088 |
| GO:0055085 | transmembrane transport | 283 | 30 | 21.48 | 0.034 | 0.092 |
| GO:0048518 | positive regulation of biological proces... | 17 | 4 | 1.29 | 0.035 | 0.093 |
| GO:0016043 | cellular component organization | 162 | 19 | 12.3 | 0.035 | 0.093 |
| GO:0051186 | cofactor metabolic process | 42 | 7 | 3.19 | 0.036 | 0.094 |
| GO:0006753 | nucleoside phosphate metabolic process | 72 | 10 | 5.47 | 0.043 | 0.109 |
| GO:0006812 | cation transport | 92 | 12 | 6.98 | 0.043 | 0.109 |
| GO:0030029 | actin filament-based process | 11 | 3 | 0.84 | 0.045 | 0.109 |
| GO:0030036 | actin cytoskeleton organization | 11 | 3 | 0.84 | 0.045 | 0.109 |
| GO:0019725 | cellular homeostasis | 35 | 6 | 2.66 | 0.045 | 0.109 |
| GO:0044257 | cellular protein catabolic process | 35 | 6 | 2.66 | 0.045 | 0.109 |
| GO:0051603 | proteolysis involved in cellular protein... | 35 | 6 | 2.66 | 0.045 | 0.109 |
| GO:0071840 | cellular component organization or bioge... | 190 | 21 | 14.42 | 0.049 | 0.116 |

**Table S5:** Genes with strongest increase in connectivity in module 27.

|  | *C. obscurior*<br>gene | *D. melanogaster*<br>ortholog | pfam domains | putative function |
| --- | --- | --- | --- | --- |
| 1 | Cobs_15828 | Protein lifeguard 4<br>(Q9DA39) | Bax1-I | Anti-apoptotic protein<br>aka Golgi anti-apoptotic<br>protein (GAAP) |
| 2 | Cobs_06161 | FBgn0011284 | RS4NT, S4,<br>Ribosomal_S4e,<br>KOW, 40S.S4_C | Ribosomal protein S4 |
| 3 | Cobs_09326 | FBgn0015756 | Ribosomal_L6 | Ribosomal protein L9 |
| 4 | Cobs_18136 | FBgn0014391 | ATP-synt_Eps | ATP synthase epsilon chain |
| 5 | Cobs_12509 | FBgn0261596 | Ribosomal_S24e | Ribosomal protein S24 |
| 6 | Cobs_17251 | Short/branched chain specific acyl-CoA<br>dehydrogenase, mitochondrial (P45954) | Acyl-CoA_dh_N, Acyl-CoA_dh_M,<br>Acyl-CoA_dh_1, Linker_histone,<br>adh_short | short/branched chain specific<br>acyl-CoA dehydrogenase,<br>mitochondrial-like |
| 7 | Cobs_17813 | FBgn0015031 | COX6C | cytochrome c oxidase subunit VIc |
| 8 | Cobs_07556 | FBgn0031059 | NA | uncharacterised |
| 9 | Cobs_07129 | 60S ribosomal protein L35<br>(Q3MHM7) | Ribosomal_L29 | 60S ribosomal protein L35 |
| 10 | Cobs_00057 | 26S proteasome complex subunit<br>SEM1 (P60897) | DSS1_SEM1 | 26S proteasome complex<br>subunit DSS1 |

## References

Aamodt, R. M. (2009). Age-and caste-dependent decrease in expression of genes maintaining dna and rna quality and mitochondrial integrity in the honeybee wing muscle. Experimental Gerontology, 44(9):586– 593.

Alexa, A. and Rahnenfuhrer, J. (2018). topGO: Enrichment Analysis for Gene Ontology. R package version 2.34.0.

Altschul, S. F., Gish, W., Miller, W., Myers, E. W., and Lipman, D. J. (1990). Basic local alignment search tool. Journal of molecular biology, 215(3):403–410.

Anders, S., Pyl, P. T., and Huber, W. (2015). Htseq—a python framework to work with high-throughput sequencing data. Bioinformatics, 31(2):166–169.

Araújo, A. R., Reis, M., Rocha, H., Aguiar, B., Morales-Hojas, R., Macedo-Ribeiro, S., Fonseca, N. A., Reboiro-Jato, D., Reboiro-Jato, M., Fdez-Riverola, F., et al. (2013). The drosophila melanogaster methuselah gene: a novel gene with ancient functions. PloS one, 8(5):e63747.

Aurori, C. M., Buttstedt, A., Dezmirean, D. S., Mărghitaş, L. A., Moritz, R. F., and Erler, S. (2014). What is the main driver of ageing in long-lived winter honeybees: antioxidant enzymes, innate immunity, or vitellogenin? Journals of Gerontology Series A: Biomedical Sciences and Medical es, 69(6):633–639.

Balistreri, C. R., Madonna, R., Melino, G., and Caruso, C. (2016). The emerging role of notch pathway in ageing: focus on the related mechanisms in age-related diseases. Ageing research reviews, 29:50–65.

Benjamini, Y. and Hochberg, Y. (1995). Controlling the false discovery rate: a practical and powerful approach to multiple testing. Journal of the Royal statistical society: series B (Methodological), 57(1):289–300.

Bielig, H., Zurek, B., Kutsch, A., Menning, M., Philpott, D., Sansonetti, P., and Kufer, T. (2009). A function for aamp in nod2-mediated nf-κb activation. Molecular immunology, 46(13):2647–2654.

Bonini, N. M., Leiserson, W. M., and Benzer, S. (1998). Multiple roles of theeyes absentgene indrosophila. Developmental biology, 196(1):42–57.

Burstein, E., Hoberg, J. E., Wilkinson, A. S., Rumble, J. M., Csomos, R. A., Komarck, C. M., Maine, G. N., Wilkinson, J. C., Mayo, M. W., and Duckett, C. S. (2005). Commd proteins, a novel family of structural and functional homologs of murr1. Journal of Biological Chemistry, 280(23):22222–22232.

Calderwood, S. K., Murshid, A., and Prince, T. (2009). The shock of aging: molecular chaperones and the he shock response in longevity and aging–a mini-review. Gerontology, 55(5):550–558.

Camacho, C., Coulouris, G., Avagyan, V., Ma, N., Papadopoulos, J., Bealer, K., and Madden, T. L. (2009). Blast+: architecture and applications. BMC bioinformatics, 10(1):421.

Carlson, M. R., Zhang, B., Fang, Z., Mischel, P. S., Horvath, S., and Nelson, S. F. (2006). Gene connectivity, function, and sequence conservation: predictions from modular yeast co-expression networks. BMC genomics, 7(1):40.

Consortium, D.. G. et al.. (2007). Evolution of genes and genomes on the drosophila phylogeny. Nate, 450(7167):203.

Consortium, U. (2018). Uniprot: a worldwide hub of protein knowledge. Nucleic acids research, 47(D1):D506–D515.

Corona, M., Hughes, K. A., Weaver, D. B., and Robinson, G. E. (2005). Gene expression patterns associated with queen honey bee longevity. Mechanisms of ageing and development, 126(11):1230– 1238.

Corona, M., Velarde, R. A., Remolina, S., Moran-Lauter, A., Wang, Y., Hughes, K. A., and Robinson, G. E. (2007). Vitellogenin, juvenile hormone, insulin signaling, and queen honey bee longevity. Proceedings of the National Academy of Sciences, 104(17):7128–7133.

Dalle-Donne, I., Rossi, R., Colombo, G., Giustarini, D., and Milzani, A. (2009). Protein s-glutathionylation: a regulatory device from bacteria to humans. Trends in biochemical sciences, 34(2):85–96.

Drummond, D. A., Bloom, J. D., Adami, C., Wilke, C. O., and Arnold, F. H. (2005). Why highly expressed proteins evolve slowly. Proceedings of the National Academy of Sciences, 102(40):14338– 14343.

Elsik, C. G., Tayal, A., Diesh, C. M., Unni, D. R., Emery, M. L., Nguyen, H. N., and Hagen, D. E. (2015). Hymenoptera genome database: integrating genome annotations in hymenopteramine. Nucleic acids research, 44(D1):D793–D800.

Elsner, D., Meusemann, K., and Korb, J. (2018). Longevity and transposon defense, the case of termite reproductives. Proceedings of the National Academy of Sciences, 115(21):5504–5509.

Emms, D. M. and Kelly, S. (2015). Orthofinder: solving fundamental biases in whole genome comparisons dramatically improves orthogroup inference accuracy. Genome biology, 16(1):157.

Fabian, D. K., Garschall, K., Klepsatel, P., Kapun, M., Lemaitre, B., Schl, C., Arking, R., and Flatt, T. (2018). Evolution of longevity improves immunity in Drosophila. Evolution Letters, 00(0):1–13.

Flatt, T. and Partridge, L. (2018). Horizons in the evolution of aging. BMC biology, 16(1):1–13.

Frenk, S. and Houseley, J. (2018). Gene expression hallmarks of cellular ageing. Biogerontology, 19(6):547– 566.

Friedrich, M. and Jones, J. W. (2016). Gene ages, nomenclatures, and functional diversification of the methuselah/methuselah-like gpcr family in drosophila and tribolium. Journal of Experimental Zoology Part B: Molecular and Developmental Evolution, 326(8):453–463.

Gems, D. and Partridge, L. (2013). Genetics of longevity in model organisms: debates and paradigm shifts. Annual review of physiology, 75:621–644.

Hartke, J., Schell, T., Jongepier, E., Schmidt, H., Sprenger, P. P., Paule, J., Bornberg-Bauer, E., Schmitt, T., Menzel, F., Pfenninger, M., et al. (2019). Hybrid genome assembly of a neotropical mutualistic ant. Genome biology and evolution, 11(8):2306–2311.

Heinze, J. and Schrempf, A. (2012). Terminal investment: individual reproduction of ant queens increases with age. PLoS One, 7(4).

Honda, Y., Tanaka, M., and Honda, S. (2010). Trehalose extends longevity in the nematode caenorhabditis elegans. Aging cell, 9(4):558–569.

Jia, K., Cui, C., Gao, Y., Zhou, Y., and Cui, Q. (2018). An analysis of aging-related genes derived from the genotype-tissue expression project (gtex). Cell death discovery, 4(1):1–14.

Jones, O. R., Scheuerlein, A., Salguero-Gómez, R., Camarda, C. G., Schaible, R., Casper, B. B., Dahlgren, J. P., Ehrlén, J., García, M. B., Menges, E. S., et al. (2014). Diversity of ageing across the tree of life. Nature, 505(7482):169.

Kanamori, Y., Saito, A., Hagiwara-Komoda, Y., Tanaka, D., Mitsumasu, K., Kikuta, S., Watanabe, M., Cornette, R., Kikawada, T., and Okuda, T. (2010). The trehalose transporter 1 gene sequence is conserved in insects and encodes proteins with different kinetic properties involved in trehalose import into peripheral tissues. Insect biochemistry and molecular biology, 40(1):30–37.

Keller, L. and Genoud, M. (1997). Extraordinary lifespans in ants : a test of evolutionary theories of ageing. Nature, 389(October):3–5.

Kenyon, C. J. (2010). The genetics of ageing. Nature, 464(7288):504.

Kim, D., Paggi, J. M., Park, C., Bennett, C., and Salzberg, S. L. (2019). Graph-based genome alignment and genotyping with hisat2 and hisat-genotype. Nature biotechnology, 37(8):907–915.

Kirkwood, T. B. (1977). Evolution of ageing. Nature, 270(5635):301–304.

Kirkwood, T. B. and Austad, S. N. (2000). Why do we age? Nature, 408(6809):233.

Kleino, A., Myllymaki, H., Kallio, J., Vanha-aho, L.-M., Oksanen, K., Ulvila, J., Hultmark, D., Valanne, S., and Ramet, M. (2008). Pirk Is a Negative Regulator of the Drosophila Imd Pathway. The Journal of Immunology, 180(8):5413–5422.

Koga, H., Kaushik, S., and Cuervo, A. M. (2011). Protein homeostasis and aging: The importance of exquisite quality control. Ageing research reviews, 10(2):205–215.

Kramer, B. H., Schrempf, A., Scheuerlein, A., and Heinze, J. (2015). Ant colonies do not trade-off reproduction against maintenance. PLoS One, 10(9):e0137969.

Kruegel, U., Robison, B., Dange, T., Kahlert, G., Delaney, J. R., Kotireddy, S., Tsuchiya, M., Tsuchiyama, S., Murakami, C. J., Schleit, J., et al. (2011). Elevated proteasome capacity extends replicative lifespan in saccharomyces cerevisiae. PLoS genetics, 7(9):e1002253.

Kuhn, J. M. M., Meusemann, K., and Korb, J. (2019). Long live the queen, the king and the commoner? transcript expression differences between old and young in the termite cryptotermes secundus. PloS one, 14(2):e0210371.

Langfelder, P. and Horvath, S. (2008). Wgcna: an r package for weighted correlation network analysis. BMC bioinformatics, 9(1):559.

Langfelder, P., Luo, R., Oldham, M. C., and Horvath, S. (2011). Is my network module preserved and reproducible? PLoS Comput Biol, 7(1):e1001057.

Laurindo, F. R., Pescatore, L. A., and de Castro Fernandes, D. (2012). Protein disulfide isomerase in redox cell signaling and homeostasis. Free Radical Biology and Medicine, 52(9):1954–1969.

Lee, B.-H., Lee, M. J., Park, S., Oh, D.-C., Elsasser, S., Chen, P.-C., Gartner, C., Dimova, N., Hanna, J., Gygi, S. P., et al. (2010). Enhancement of proteasome activity by a small-molecule inhibitor of usp14. Nature, 467(7312):179.

Li, H., Handsaker, B., Wysoker, A., Fennell, T., Ruan, J., Homer, N., Marth, G., Abecasis, G., and Durbin, R. (2009). The sequence alignment/map format and samtools. Bioinformatics, 25(16):2078–2079.

Lin, Y.-J., Seroude, L., and Benzer, S. (1998). Extended life-span and stress resistance in the drosophila mutant methuselah. Science, 282(5390):943–946.

Loch, G., Zinke, I., Mori, T., Carrera, P., Schroer, J., Takeyama, H., and Hoch, M. (2017). Antimicrobial peptides extend lifespan in Drosophila. PLoS ONE, 12(5):1–15.

Lockett, G. A., Almond, E. J., Huggins, T. J., Parker, J. D., and Bourke, A. F. (2016). Gene expression differences in relation to age and social environment in queen and worker bumble bees. Experimental gerontology, 77:52–61.

López-Otín, C., Blasco, M. A., Partridge, L., Serrano, M., and Kroemer, G. (2013). The hallmarks of aging. Cell, 153(6):1194–1217.

Love, M. I., Huber, W., and Anders, S. (2014). Moderated estimation of fold change and dispersion for rna-seq data with deseq2. Genome biology, 15(12):550.

Löytynoja, A. (2014). Phylogeny-aware alignment with prank. In Multiple sequence alignment methods, pages 155–170. Springer.

Lucas, E. R. and Keller, L. (2018). Elevated expression of ageing and immunity genes in queens of the black garden ant. Experimental gerontology, 108:92–98.

Lucas, E. R., Privman, E., and Keller, L. (2016). Higher expression of somatic repair genes in long-lived ant queens than workers. Aging (Albany NY), 8(9):1940.

Lucas, E. R., Romiguier, J., and Keller, L. (2017). Gene expression is more strongly influenced by age than caste in the ant lasius niger. Molecular ecology, 26(19):5058–5073.

Mistry, J., Bateman, A., and Finn, R. D. (2007). Predicting active site residue annotations in the pfam database. BMC bioinformatics, 8(1):298.

Mitchell, A., Chang, H.-Y., Daugherty, L., Fraser, M., Hunter, S., Lopez, R., McAnulla, C., McMenamin, C., Nuka, G., Pesseat, S., et al. (2014). The interpro protein families database: the classification resource after 15 years. Nucleic acids research, 43(D1):D213–D221.

Negroni, M. A., Foitzik, S., and Feldmeyer, B. (2019). Long-lived temnothorax ant queens switch from investment in immunity to antioxidant production with age. Scientific reports, 9(1):7270.

Negroni, M. A., Jongepier, E., Feldmeyer, B., Kramer, B. H., and Foitzik, S. (2016). Life history evolution in social insects: a female perspective. Current opinion in insect science, 16:51–57.

Oettler, J. and Schrempf, A. (2016). Fitness and aging in cardiocondyla obscurior ant queens. Current opinion in insect science, 16:58–63.

Partridge, L., Alic, N., Bjedov, I., and Piper, M. D. (2011). Ageing in drosophila: the role of the insulin/igf and tor signalling network. Experimental gerontology, 46(5):376–381.

R Core Team (2018). R: A Language and Environment for Statistical Computing. R Foundation for Statistical Computing, Vienna, Austria.

Rivera, M. C., Jain, R., Moore, J. E., and Lake, J. A. (1998). Genomic evidence for two functionally distinct gene classes. Proceedings of the National Academy of Sciences, 95(11):6239–6244.

Rubinsztein, D. C., Mariño, G., and Kroemer, G. (2011). Autophagy and aging. Cell, 146(5):682–695.

Schrader, L., Kim, J. W., Ence, D., Zimin, A., Klein, A., Wyschetzki, K., Weichselgartner, T., Kemena, C., Stökl, J., Schultner, E., et al. (2014). Transposable element islands facilitate adaptation to novel environments in an invasive species. Nature communications, 5(1):1–10.

Schrempf, A., Heinze, J., and Cremer, S. (2005). Sexual cooperation: mating increases longevity in ant queens. Current Biology, 15(3):267–270.

Seehuus, S.-C., Taylor, S., Petersen, K., and Aamodt, R. M. (2013). Somatic maintenance resources in the honeybee worker fat body are distributed to withstand the most life-threatening challenges at each life stage. PloS one, 8(8):e69870.

Shannon, P., Markiel, A., Ozier, O., Baliga, N. S., Wang, J. T., Ramage, D., Amin, N., Schwikowski, B., and Ideker, T. (2003). Cytoscape: a software environment for integrated models of biomolecular interaction networks. Genome research, 13(11):2498–2504.

Southworth, L. K., Owen, A. B., and Kim, S. K. (2009). Aging mice show a decreasing correlation of gene expression within genetic modules. PLoS genetics, 5(12):e1000776.

Suyama, M., Torrents, D., and Bork, P. (2006). Pal2nal: robust conversion of protein sequence alignments into the corresponding codon alignments. Nucleic acids research, 34(Suppl 2):W609–W612.

Tatar, M., Bartke, A., and Antebi, A. (2003). The endocrine regulation of aging by insulin-like signals. Science, 299(5611):1346–1351.

Thurmond, J., Goodman, J. L., Strelets, V. B., Attrill, H., Gramates, L. S., Marygold, S. J., Matthews, B. B., Millburn, G., Antonazzo, G., Trovisco, V., et al. (2018). Flybase 2.0: the next generation. Nucleic acids research, 47(D1):D759–D765.

Tilstra, J. S., Robinson, A. R., Wang, J., Gregg, S. Q., Clauson, C. L., Reay, D. P., Nasto, L. A., St Croix, C. M., Usas, A., Vo, N., et al. (2012). Nf-κb inhibition delays dna damage–induced senescence and aging in mice. The Journal of clinical investigation, 122(7):2601–2612.

Tomaru, U., Takahashi, S., Ishizu, A., Miyatake, Y., Gohda, A., Suzuki, S., Ono, A., Ohara, J., Baba, T., Murata, S., et al. (2012). Decreased proteasomal activity causes age-related phenotypes and promotes the development of metabolic abnormalities. The American journal of pathology, 180(3):963–972.

Turan, Z. G., Parvizi, P., Dönertaş, H. M., Tung, J., Khaitovich, P., and Somel, M. (2019). Molecular footprint of medawar’s mutation accumulation process in mammalian aging. Aging cell, 18(4):e12965.

Vilchez, D., Morantte, I., Liu, Z., Douglas, P. M., Merkwirth, C., Rodrigues, A. P., Manning, G., and Dillin, A. (2012). Rpn-6 determines c. elegans longevity under proteotoxic stress conditions. Nature, 489(7415):263.

Von Wyschetzki, K., Rueppell, O., Oettler, J., and Heinze, J. (2015). Transcriptomic signatures mirror the lack of the fecundity/longevity trade-off in ant queens. Molecular biology and evolution, 32(12):3173– 3185.

Ward, P. S., Brady, S. G., Fisher, B. L., and Schultz, T. R. (2015). The evolution of myrmicine ants: phylogeny and biogeography of a hyperdiverse ant clade (h ymenoptera: F ormicidae). Systematic Entomology, 40(1):61–81.

Williams, G. C. (1957). Pleiotropy, natural selection, and the evolution of senescence. evolution, pages 398–411.

Yang, Z. (1997). Paml: a program package for phylogenetic analysis by maximum likelihood. Bioinformatics, 13(5):555–556.

